# Relative strength variability measures for brain structural connectomes and their relationship with cognitive functioning

**DOI:** 10.1101/2025.03.15.643458

**Authors:** Hon Wah Yeung, Colin R. Buchanan, Joanna Moodie, Ian J. Deary, Elliot M. Tucker-Drob, Mark E. Bastin, Heather C. Whalley, Keith M. Smith, Simon R. Cox

**Affiliations:** Lothian Birth Cohorts, Department of Psychology, University of Edinburgh, Edinburgh, United Kingdom; Department of Psychiatry, University of Edinburgh, Edinburgh, United Kingdom; Department of Psychology, University of Texas, Austin, TX, USA; Population Research Center and Center on Aging and Population Sciences, University of Texas at Austin, TX, USA; Centre for Clinical Brain Science, University of Edinburgh, Edinburgh, United Kingdom; Department of Computer and Information Sciences, University of Strathclyde, Glasgow, United Kingdom

**Author notes:** These authors share joint senior authorship (Keith M. Smith, Simon R. Cox).

## Abstract

In this work, we propose a new class of graph measures for weighted connectivity information in the human brain based on node relative strengths: relative strength variability (RSV), measuring susceptibility to targeted attacks, and hierarchical RSV (hRSV), a first weighted statistical complexity measure for networks. Using six different network weights for structural connectomes from the UK Biobank, we conduct comprehensive analyses to explore relationships between the RSV and hRSV, and (i) other known network measures, (ii) general cognitive function (‘*g*’). Both measures exhibit low correlations with other graph measures across all connectivity weightings indicating that they capture new information of the brain connectome. We found higher *g* was associated with lower RSV and lower hRSV. That is, higher *g* was associated with higher resistance to targeted attack and lower statistical complexity. Moreover, the proposed measures had consistently stronger associations with *g* than other widely used graph measures including clustering coefficient and global efficiency and were incrementally significant for predicting *g* above other measures for five of the six network weights. Overall, we present a new class of weighted network measures based on variations of relative node strengths which significantly improved prediction of general cognition from traditional weighted structural connectomes.

## Introduction

Decline of cognitive skills in the older population is associated with a decrease in the ability to perform necessary activities for independent living, resulting in poorer quality of life and a substantial burden on both the individual and society (Deary et al., 2009; Eshkoor et al., 2015). Understanding the neurobiological bases of cognitive aging is important for developing interventions to reduce this rate of decline. The advent of diffusion magnetic resonance imaging (dMRI) methods has allowed researchers to quantify aspects of the microstructural environment of white matter in vivo and to interrogate how individual differences in these brain properties relate to differences in cognitive functioning (Kennedy and Raz, 2009). It is clear that there are modest but consistent associations between white matter dMRI properties and cognitive functioning in ageing, though common approaches are mainly restricted to voxels only in skeletonised white matter which are well-aligned across subjects, such as Tract-Based Spatial Statistics (TBSS; Smith et al. (2006)), or as averages across major white matter pathways (Damoiseaux et al., 2009; Mayo et al., 2019; Penke et al., 2010). Advances in dMRI and neuroimaging have enabled the measurement of the structural connectome to map many hundreds of white matter connections between distal grey matter regions at the network level (Sporns et al., 2005). The connectome approach allows the characterisation of these connections using graph-theory measures, offering unique biologically tractable opportunities to understand the organising principles of the brain, and cerebral bases of cognitive differences in aging (Blesa et al., 2021b; Li et al., 2020; Smith et al., 2019; Van Den Heuvel and Sporns, 2011; Yeung et al., 2022).

The human structural connectome has a rich-club of strongly interconnected hub regions (Van Den Heuvel and Sporns, 2011). More recently, the hierarchical structure of the binarised connectome has been expanded using computational modelling to a four-tier structure, with Tier 1 being this cognitive processing rich-club, Tier 2 involving mostly regions involved in sensorimotor processing, Tier 3 involving heteromodal regions, and Tier 4 linked to memory and emotion (Blesa et al., 2021a; Smith et al., 2019). Here, we propose novel weighted measures for analysing the hierarchical organisation of more information-rich weighted connectomes. This is important because connectivity weights contain biologically relevant information about the microstructural environment which is pertinent to maintaining brain function, such as axonal myelination; (Beaulieu, 2002; Jones et al., 2013). Particularly, we study relative node strengths and present two measures for analysing their global variability: network-wide variation, which we call relative strength variability (RSV), and variations across narrow bands, or tiers, of the weighted node hierarchy using a sliding window approach (hRSV). The former is a measure which is sensitive to the strength of connectivity between high and low degree nodes, i.e. strength of connectivity between the network core and periphery. We explain how the latter is akin to a weighted measure of hierarchical complexity, capturing the diversity of connectivity patterns in the network across similarly weighted nodes. Figure 1 A depicts the types of networks which have high and low values of RSV and hRSV.

**Figure 1:**
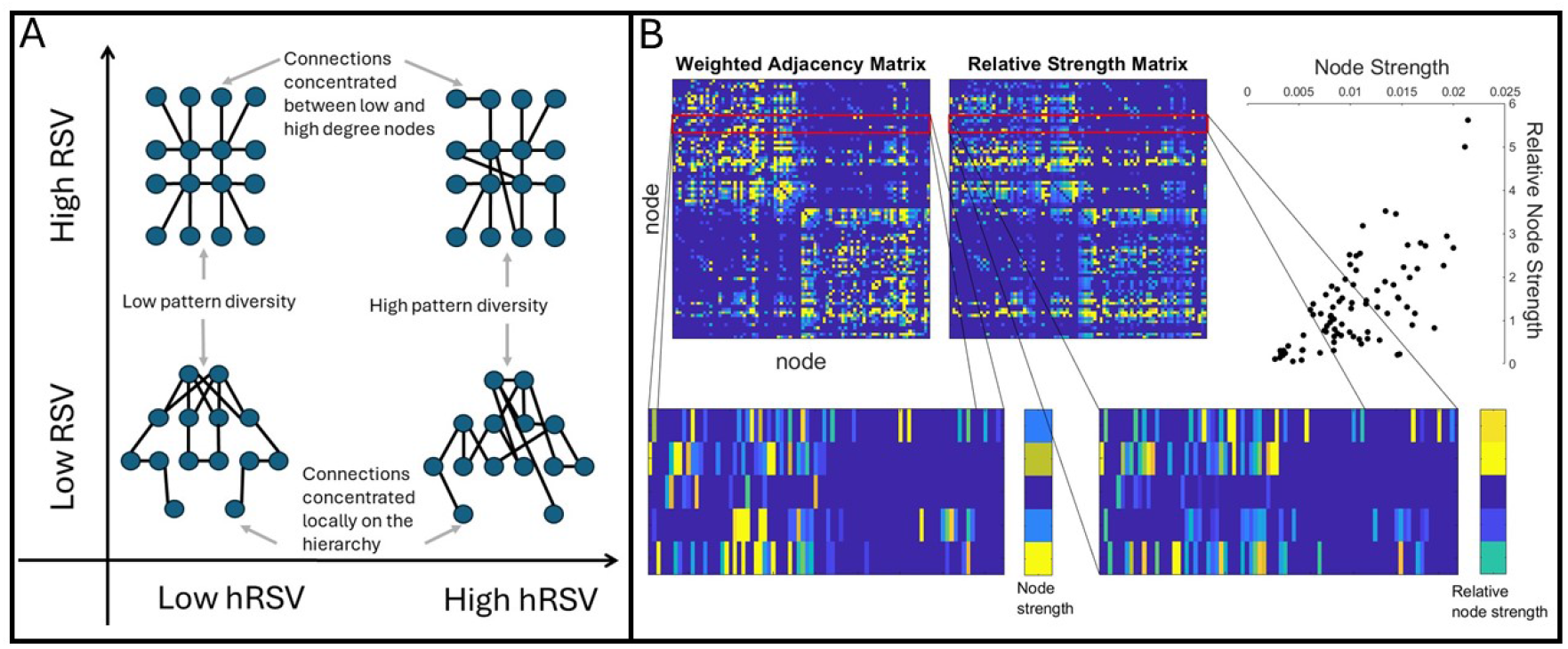
A. Illustration of the connectivity profiles captured by the proposed measures, Relative Strength Variability (RSV) and hierarchical Relative Strength Variability (hRSV). For ease of conceptualisation, these are provided for binary networks using the degree as an analogy to node strength. B. Illustrates the difference between node strengths and relative node strengths. Nodes which generally connect to stronger nodes will have high relative node strength, while nodes which connect to lower strength nodes will have low relative node strength. The relationship between node strength and relative node strength is moderately correlated in this example, as shown in the scatter plot, top right.

Weighted graph theory analysis of brain connectomic data is however not without its challenges. There are typically high correlations among many commonly-used graph theory measures in weighted connectomes which may also partly differ as a function of user-defined thresholding approaches (Madole et al., 2023). Clearly it would be preferable not to lose the rich information contained in the connection weights, but novel measures of brain connectivity should i) offer a biologically tractable description of the brain and its functioning, ii) be related to differences in cognitive function, and iii) ensure it does not simply recapitulate information already available via more conventional measures. To this end, we subject our newly proposed measures to a comprehensive set of comparisons with other weighted network measures (node strength variance, assortativity, normalised clustering coefficient, routing efficiency, diffusion efficiency) across six connectivity weightings, fractional anisotropy (FA), mean diffusivity (MD), intracellular volume fraction (ICVF), isotropic volume fraction (ISOVF) and orientation dispersion (OD), using UK Biobank (UKB) connectome data.

Prior work indicated that there are high correlations among many graph theory measures when applied to the human brain connectome (Madole et al., 2023). However, since the two new variability measures (RSV and hRSV) were derived from a distinct statistical and theoretical framework, we hypothesised that they would be only moderately correlated with common graph measures. The cerebral correlates of *g* are widely distributed throughout the brain (Barbey et al., 2012; Camilleri et al., 2018; Cox et al., 2019; Gläscher et al., 2010) and higher score in the RSV and hRSV represent a lower overall resistent to targeted attack. We therefore hypothesised that higher RSV and hRSV scores would be associated with lower *g* scores, and would account for unique variance in *g* beyond other graph measures. We examine i) their relationships with other brain measures and common graph metrics, ii) their associations with general cognitive function (*g*) and iii) whether the novel measures account for incrementally significant variance in *g* beyond the structural brain measures and graph metrics that have been conventionally studied.

## Materials and Methods

### Relative Strength Variability Measures

The motivation for the proposed measures originates in formulating a weighted measure of hierarchical complexity. Hierarchical complexity in binary networks assesses the diversity of connectivity patterns across the degree hierarchy of the network (Smith et al., 2019). It has recently been shown to be a measure of statistical complexity in networks, which means it is low for both regular and random networks, but high for structurally diverse networks such as real-world networks (Smith and Smith, 2024). It is defined using variances of the neighbourhood degree sequences across nodes of the same degree, where the neighbourhood degree sequence of node *i* is the sorted list of degrees of nodes to which *i* is connected.

Here we introduce two separate but related strength variability measures. The first – RSV – aims to assess the variance in relative node strength patterns across the whole network. To calculate this, we first compute the relative strength of a node compared to the nodes it is connected to. Figure 1B illustrates the relationship between node strength and relative node strength in an example weighted connectome. We then take the standard deviation of these measurements over all nodes. The highest values for this measure should come from networks which connect to nodes of dissimilar connectivity strength. Because the most dissimilar connectivity strengths will be between highest and lowest node strengths in the network, this measurement is particularly sensitive to differences in core-periphery connectivity. High measurements will be obtained where connectivity is concentrated between low and high strength nodes, while low measurements will be obtained where connectivity is concentrated more locally on the strength hierarchy, Figure 1A.

The second is a sliding window variant – hRSV – which, instead of computing this variability over all nodes, first defines a sliding window on node strengths, and computes the variability over all nodes whose strength lies within that window. Finally, the average over all windows from smallest to largest in node strength is taken. This assesses the diversity of connectivity patterns of similar strength within the network. Low values will be obtained by networks with either highly regular or highly random connectivity patterns, while higher values will be obtained in networks with a greater diversity of connectivity patterns, Fig 1A. Therefore, we can expect these measurements to pick up distinct information within weighted networks. Note, Fig 1A demonstrates cases of high RSV and low hRSV, high RSV and high hRSV, low RSV and low hRSV, and low RSV and high hRSV.

### Relative Strength Variability (RSV)

For densely weighted networks such as structural connectomes, we want to replace neighbourhood degree sequences (Smith et al., 2019) with something similar involving node strengths, where the node strength of node *i* is the sum of edges’ weights adjacent to 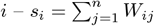 where *W* is the weighted adjacency matrix. We then define the set of nodes strengths in the network as 𝒮 = {*s*_1_, *s*_2_, …, *s*_*n*_}. However, in this case, we cannot expect node strengths to ever become equal in the same way that degrees can be, and so we want to instead consider how to quantify the relative connectivity profile of each node as a single feature and then assess the variability of those connectivity profiles across windows of connectivity strength.

To do this, we consider the relative node strength of a node, with respect to its neighbours. We therefore define the relative node strength of node *i* as the node strength of *i* divided by the mean of the node strengths of its neighbours:

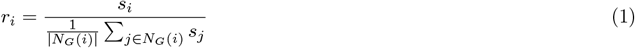

where *N*_*G*_(*i*) is the set of neighbourhood nodes of *i*. An illustration of relative node strength compared to node strength can be seen in figure 1 B.

This dramatically simplifies the measurement of hierarchical complexity as defined for binary networks, since now we are no longer strictly confined to assessing variances across nodes of the same degree, but can assess variances across any set of nodes in the network.

To this end, the simplest set to consider is just the entire network. We therefore define the Relative Strength Variability as:

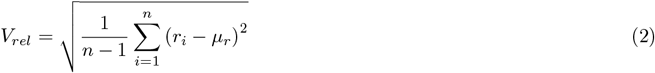

where

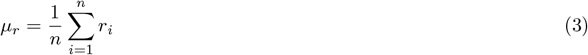

This measures the extent to which the connections in the network are connected to similar strength nodes. It is high for networks with many connections between high and low degree nodes, star-like networks, and low for networks whose connections are concentrated to networks with similar strengths, with a more localised hierarchical structure, as illustrated by the *y*-axis in Figure 1 A.

### Sliding window variant (hierarchical RSV; hRSV)

We formulate a sliding window variant (hRSV) by taking the average of the standard deviations of relative strengths of subsets of nodes across the node strength spectrum. Firstly, the node relative strengths are sorted according to the mean node strengths of the population. Then, we choose a window size, *w*, and take the standard deviation of the relative node strengths of all nodes of strength within *w*. We slide *w* across from the minimum strength to the maximum strength. This results in *n* − *w* + 1 standard deviations. We get the global measure by taking average over all windows.

We propose that by choosing a suitably small *w*, we obtain a weighted measure of hierarchical complexity. Hierarchical complexity should be low for ER random graphs, random geometric graphs and highly hierarchical graphs, but large in heterogeneous geometric graphs (random geometric graphs with heterogeneous degree distributions) (Smith and Smith, 2024). We demonstrate this behaviour through simulations in Supplementary Materials section A. Particularly, we demonstrate that a small window size best distinguishes the hRSV measure from the global RSV. For the real data, we chose window sizes of 4 (for small window size) and 21 (21 correspond to 1/4 of the number of nodes).

### Participants

The UKB is a large-scale epidemiology study which recruited approximately 500,000 community-dwelling, generally healthy subjects aged 40–69 years from across Great Britain between 2006 and 2010 (Sudlow et al., 2015). Participants provided comprehensive demographic, psychosocial, medical information and cognitive testing during an initial visit to a UKB assessment centre. Between 2014 and 2020 a subset of approximately 40,000 participants underwent brain MRI (Miller et al., 2016). UKB received ethical approval from the North West Multi-centre Research Ethics Committee (REC reference 11/NW/0382). The current study was conducted under approved UKB application number 10279.

### Image Processing and Structural Networks

At the time of processing, *N* = 37, 518 participants with compatible T1-weighted and dMRI data were available from UKB, and the structural connectomes for these participants were derived by the team at University of Edinburgh, as previously described in Buchanan et al. (2020) and Yeung et al. (2022), and summarised briefly below.

All imaging data were acquired using Siemens Skyra 3T scanners, with a 32-channel Siemens head radiofrequency coil, (Siemens Medical Solutions, Erlangen, Germany; see http://biobank.ctsu.ox.ac.uk/crystal/refer.cgi?id=2367) at four UK sites: Cheadle (N = 21, 827, 58%), Reading (N = 5,706, 16%), Newcastle (N = 9,700, 26%), and Bristol (N = 51, 0.13%). Details of the MRI protocol and preprocessing are freely available (Alfaro-Almagro et al., 2018; Miller et al., 2016).

3D T1-weighted volumes were acquired using a magnetization-prepared rapid gradient-echo sequence at 1 × 1 × 1 mm resolution with 208 × 256 × 256 field of view. The dMRI data were acquired using a spin-echo echo-planar imaging sequence (50 b = 1000 s/mm2, 50 b = 2000 s/mm2 and 10 b = 0 s/mm2) resulting in 100 distinct diffusion-encoding directions. The field of view was 104 × 104 mm with imaging matrix 52 × 52 and 72 slices with slice thickness of 2 mm resulting in 2 × 2 × 2 mm voxels. Each T1-weighted volume was parcellated into 85 distinct neuroanatomical Regions-Of-Interest (ROI) with FreeSurfer v6.0 and 34 cortical structures per hemisphere were identified using the Desikan-Killany atlas (Desikan et al., 2006). The brain-stem, accumbens area, amygdala, caudate nucleus, hippocampus, pallidum, putamen, thalamus and ventral diencephalon were also extracted with FreeSurfer. Whole-brain tractography was performed using a probabilistic algorithm (BEDPOSTX/ProbtrackX; (Behrens et al., 2007, 2003)), with tracking initiated from white matter voxels and using criteria as described previously (Buchanan et al., 2020). Water diffusion parameters were estimated for FA, which measures the directionality of water molecule diffusion, and for MD, which measures the magnitude of diffusion. The parameters obtained from NODDI were: ICVF which measures neurite density; ISOVF which measures extracellular water diffusion; and OD which measures the degree of fanning or angular variation in neurite orientation (Zhang et al., 2012). After aligning ROIs from T1-weighted to diffusion space, networks were constructed for each participant and represented as 85 × 85 adjacency matrices, with pairwise connections between the 85 ROIs identified by the streamlines obtained from probabilistic tractography. For each participant, six network weightings were computed. Streamline count (SC) was computed by recording the total streamline count (uncorrected) between each pair of ROIs. In addition, five further network weightings (FA, MD, ICVF, ISOVF and OD) were computed by recording the mean value of the diffusion parameter in voxels identified along all interconnecting streamlines between each pair of ROIs.

Following local quality checks of the connectome outputs and exclusions for processing errors, connectomes were obtained for 37,284 participants. Participants with missing demographic information (age, sex, MRI site and head position), missing global brain measures (white matter volume (WMV), gray matter volume (GMV) and intracranial volume (ICV)) were excluded. Participants with other major neurological disorders (for example dementia, Parkinson’s disease, stroke, multiple sclerosis and other chronic degenerative neurological problems were also excluded (Moodie et al., 2024). In total, 35, 529 participants (44.6–82.7 years of age, 46.6% male) remained after participants were excluded following. We applied proportional thresholding approach to the structural connectomes, only retaining connections that are present in at least 50% of subjects.

### Common graph measures

The RSV measures were computed for all connectomes based on each of the six network weights alongside the following five graph measures for comparison:

1. Node strength variance (Snijders, 1981):

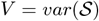

It measures the heterogeneity of the network, the spread of the node strength distribution.
2. Assortativity (Foster et al., 2010; Newman, 2002):

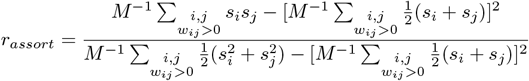

where M is the number of edges, *s*_*i*_ is the strength of node *i*. This measures the degree correlation between neighbouring nodes.
3. Normalised clustering coefficient (Onnela et al., 2005; Smith and Escudero, 2017; Watts and Strogatz, 1998):

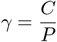

where *P* = 2*M/n*(*n* − 1), is the graph density, 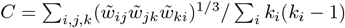 is the clustering coefficient for weighted networks, *n* is the number of nodes, *k*_*i*_ is the degree of node *i*. It measures the segregation between local clusters.
4. Routing efficiency (Goñi et al., 2013; Latora and Marchiori, 2001):

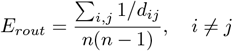

where *d*_*ij*_ is the shortest path between node *i* and *j*. It measures the global integration of the network and communication efficiency within the network.
5. Diffusion efficiency (Goñi et al., 2013; Wang and Pei, 2008):

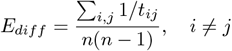

where *t*_*ij*_ is the mean first-passage time for node *i* and *j*. It complements the measure of routing efficiency.

### Other Imaging Variables

The estimation of, respectively, normal-appearing GMV and WMV, brain atrophy (ICV - total brain volume, corrected for ICV) and mean edge weights, which are imaging-derived phenotypes generated by the UKB (Alfaro-Almagro et al., 2018).

### Cognitive Tasks and *g*

The cognitive tasks included the Verbal Numerical Reasoning (VNR), Reaction Time (RT, log-transformed), Pairs Matching (PairsMatch, log (x+1) transformed), Numerical Memory (NumerMemory) Prospective Memory (ProsMemory), Trail Making Test part b (TMTb, log-transformed), Matrix pattern completion (Matrix), Symbol digit substitution (DigSym), Tower rearranging (Tower) and Paired associate learning (PairedAssociate) tests (Fawns-Ritchie and Deary, 2020; Lyall et al., 2016). Participants with missing values for more than 7 out of 10 cognitive tasks were excluded. Out of 37, 284 participants, 33, 013 remained after exclusion. Missing values were imputed via low-rank matrix recovery (Davenport and Romberg, 2016).

Principal component analysis (PCA) implemented in *pca* function in MATLAB 2022b was applied to the 10 cognitive tasks. *g* was the first unrotated principal component score (*z*-normalized), which accounted for 34.55% of the variance. The resulting *g* had a range [-4.8914, 3.5510]. The standardised loadings of scores for each task on *g* are shown in Table 2. The loadings ranged from absolute values of 0.42 to 0.75.

### Extracting unique contributions to *g*

In order to assess the incremental validity of the new measure for predicting *g*, we conducted multivariate linear regression analysis by building progressive models to estimate the unique variance in the *g* that can be explained by RSV over and above other MRI brain measures. For each combination of connectome weight and thresholding approach, we built 3 different models:

1. *g* ∼ Intercept + Covariates + RSV
2. *g* ∼ Intercept + Covariates + Mean Edge Weight + RSV
3. *g* ∼ Intercept + Covariates + Mean Edge Weight + Common Graph Measures + RSV where the covariates are age, sex and MRI site.

The formula above represents the exact column ordering of the data matrix, with the predictors sorted according to how common or conventional they are. To extract the incremental contribution in predicting *g* in the progressive models, we applied a sequential residualisation to the predictors before putting them into the linear model for predicting *g*. The sequential residualisation is equivalent to performing the Gram-Schmidt Orthogonalization process on a centered *n*-by-*p* matrix *X*, where *n* is the number of observations and *p* is the number of predictors (Björck, 1967). The pseudo-code is shown in Algorithm 1.

This sequential procedure ensures the statistical independence by removing the effects of the preceding columns, making it an appropriate candidate for studying the incremental prediction of *g*.

#### Algorithm 1

Gram-Schmidt Orthogonalisation for Data Matrix *X*

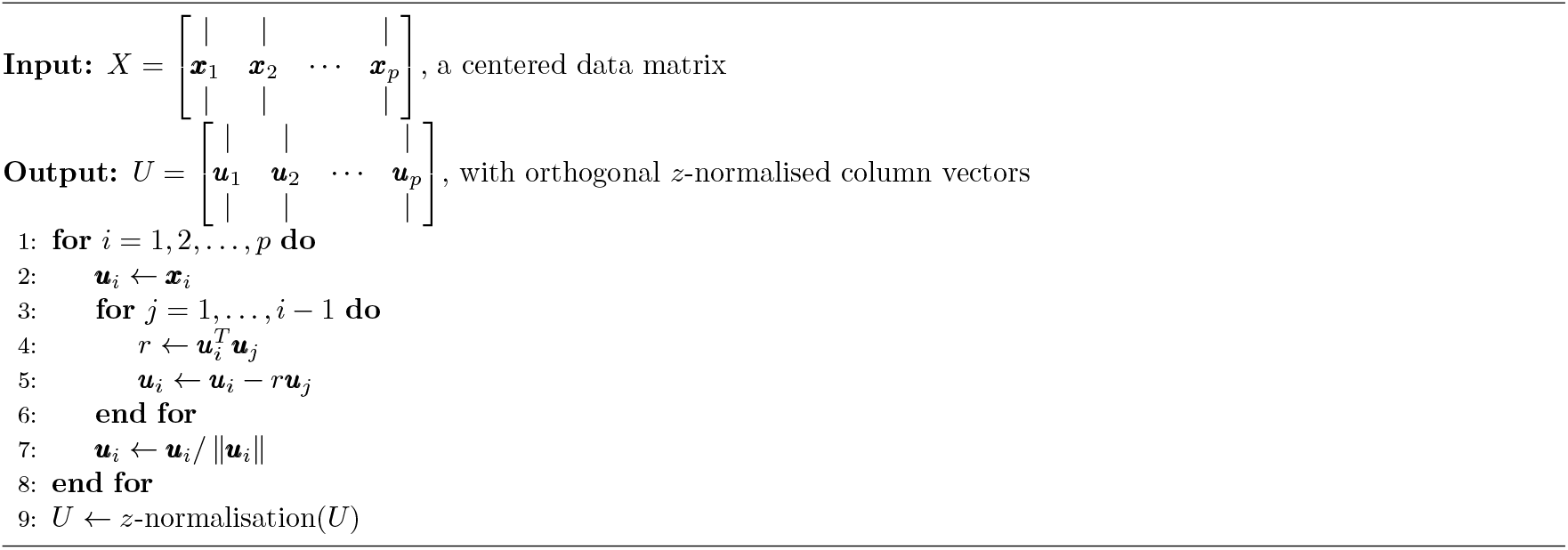

## Results

Table 1 shows participants characteristics, demographic information and cognitive task scores. Appendix I describes how we define hRSV as measure of statistical complexity for weighted networks. This is the first such measure to be proposed. Particularly, we note that taking a small enough window *w* for hRSV is important to get the correct behaviour of a statistical complexity measure and distinguish it from RSV. We now describe the main results with respect to structural connectomes and *g*.

**Table 1:** Summary characteristics of the UK Biobank participants included in the study. VNR: Verbal Numerical Reasoning. RT: Reaction Time. Pairs Match: Pairs Matching. For the prospective memory test, **1** means recall at the first attempt and **0** otherwise.

| Sample characteristics | Summary | Total Data Available (N) |
| --- | --- | --- |
| Sex [F:M] | 18,988 : 16,541 | 35,529 |
| Age (in years) [Mean (SD)] | 63.92 (7.65) | 35,529 |

(a) Demographic information for the training and test sets of the UK Biobank participants
| Cognitive measure | Sample mean score | Data Available (N) |
| --- | --- | --- |
| RT (in $\log x$) [Mean (SD)] | 6.37(0.17) | 32,972 |
| NumerMemory [Mean (SD)] | 6.76(1.27) | 24,723 |
| VNR [Mean (SD)] | 6.60 (2.04) | 32,593 |
| TMTb (in $\log x$) [Mean (SD)] | 6.28 (0.36) | 23,420 |
| Matrix [Mean (SD)] | 7.98 (2.12) | 23,977 |
| Tower [Mean (SD)] | 9.88 (3.22) | 23,775 |
| DigSym [Mean (SD)] | 18.87 (5.25) | 24,029 |
| PairedAssociate [Mean (SD)] | 6.94 (2.60) | 24,172 |
| ProsMemory [1:0] | 27,747 : 5316 | 33,063 |
| PairsMatch (in $\log(x + 1)$) [Mean (SD)] | 1.36 (0.63) | 33,078 |

(b) Sample mean scores of the cognitive tasks completed by the UK Biobank participants included in the study

**Table 2:** Standardised loadings of individual cognitive test scores on the cognitive *g*. VNR: Verbal Numerical Reasoning. RT: Reaction Time. Pairs Match: Pairs Matching. PropVar: Proportion of variance explained by *g*.

| **Cognitive Tasks** | **Loadings** |
| --- | --- |
| RT | -0.4530 |
| NumerMemory | 0.5256 |
| VNR | 0.7069 |
| TMTb | -0.7458 |
| Matrix | 0.6686 |
| Tower | 0.6122 |
| DigSym | 0.6510 |
| ProsMemory | 0.4577 |
| PairsMatch | -0.4218 |
| PropVar | 0.3451 |

### Correlations among the variables

We found that the RSV and hRSV based on binarised connectomes, MD, FA, OD and ICVF were highly correlated with each other (*r* = 0.64 to 0.97). The RSV based on ISOVF had moderate correlations with other weighted RSV (*r* = 0.28 to 0.62). SC was not highly correlated with any of the other 5 measures (*r* = −0.042 to 0.17), see Figure 2a. As for the five chosen common graph measures, the node strength variance, normalised clustering coefficient and routing efficiency were highly correlated with each other (*r* = 0.87 to 0.97), and these three measures had low-to-moderate correlations (in terms of absolute correlations) with assortativity and diffusion efficiency (|*r*| = 0.01 to 0.24). Assortativity and diffusion efficiency were moderately correlated with each other based on all connectome weights except for SC (|*r*| = 0.33 to 0.41). Figure 3 shows the mean correlation among the graph measures across five different network weights except for SC. The RSV generally showed the strongest correlations with diffusion efficiency (|*r*| = 0.31 − 0.68). It had a weak-to-moderate correlation with strength variance, clustering coefficient and routing efficiency (|*r*| = 0.005 to 0.28). Whereas for SC, we see that the RSV and hRSV were strongly negatively correlated with clustering coefficients, strength variance and routing efficiency, see Figure A.3 in the Supplementary Materials A.1. The RSV also had weak-to-moderate correlations with mean edge weights, WMV, GMV and atrophy (|*r*| : range = 0.002 to 0.28, mean = 0.08), see Figure 2a.

**Figure 2:**
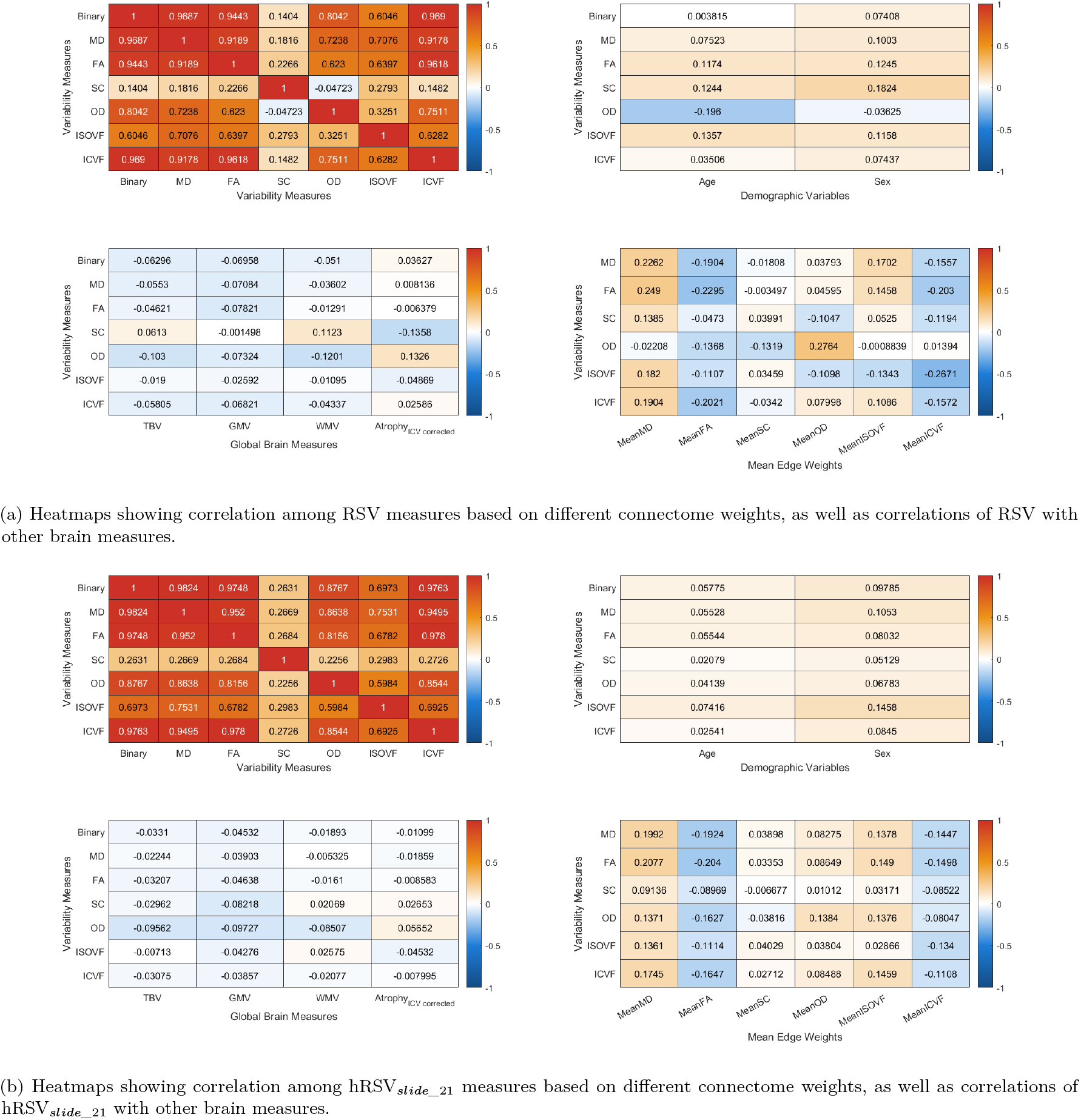
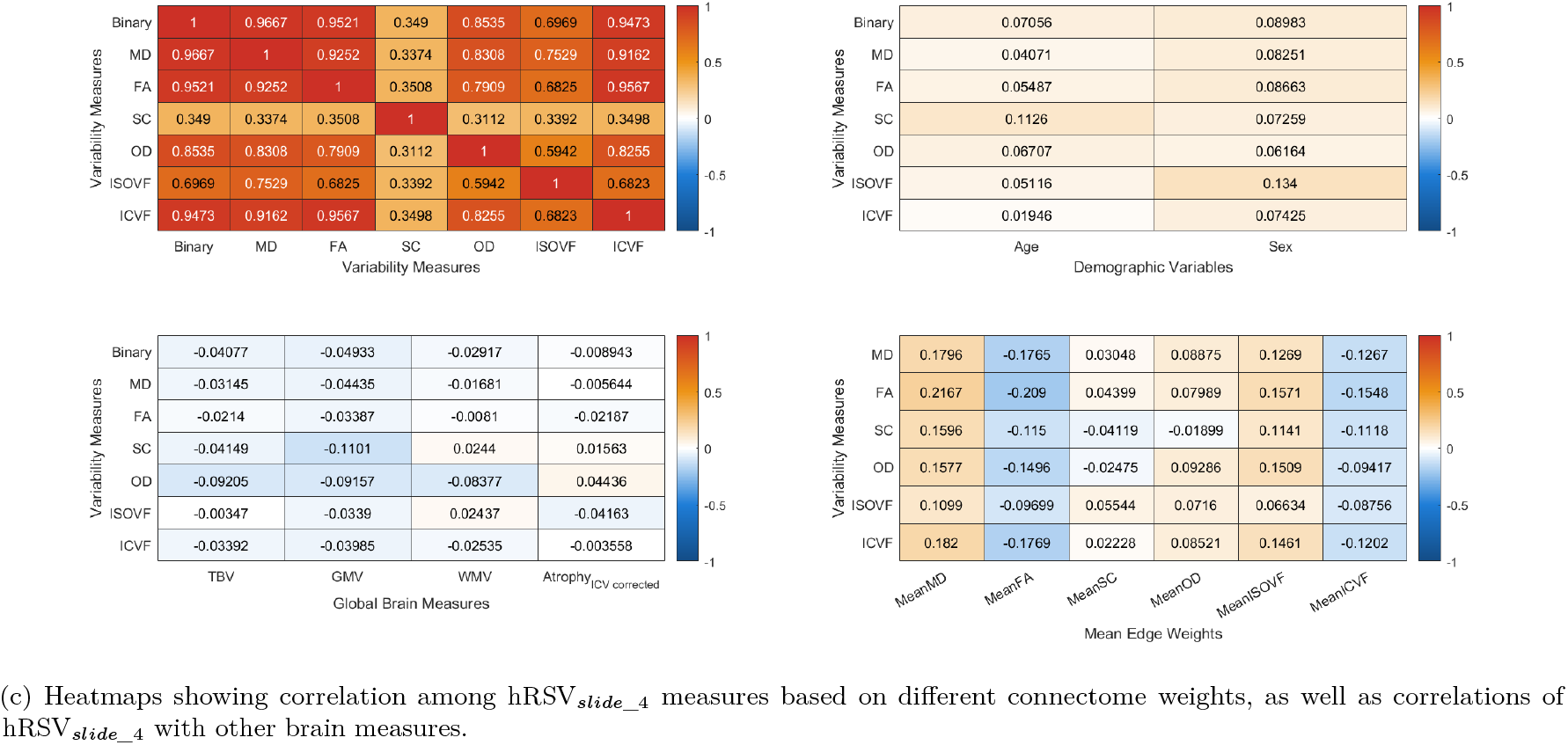
Heatmaps showing correlation among variability measures based on different connectome weights, as well as correlations of variability with other brain measures. Binary = binarised connectome, MD = mean diffusivity, FA = fractional anisotropy, SC = streamline count, OD = orientation dispersion, ISOVF = isotropic volume fraction, ICVF = intra-cellular volume fraction. TBV = Total Brain Volume, Atrophy = Brain Atrophy Volume, GMV = Gray Matter Volume, WMV = White Matter Volume

**Figure 3:**
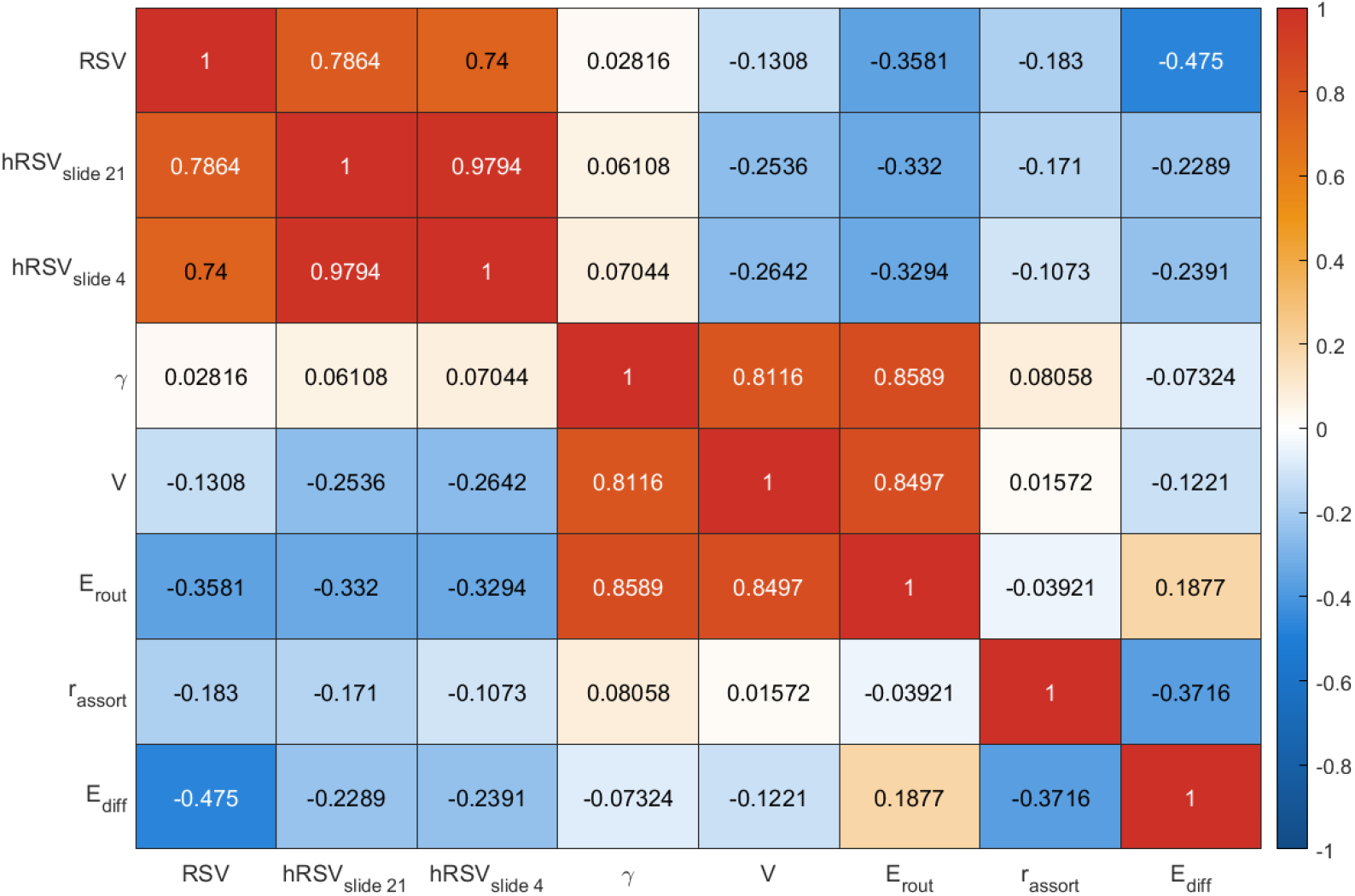
Heatmaps showing correlation among the five chosen graph metrics based on different connectome weights. RSV = Relative Strength Variability, hRSV_*slide*_*n*_ = Relative Strength Variability sliding window variant where *n* is the window size, *γ* = Clustering Coefficient, *r*_*assort*_ = assortativity, *V* = node Strength Variance, *E*_*diff*_ = Global Diffusion Efficiency, *E*_*rout*_ = Global Routing Efficiency

We found that the weighted hRSV based on binarised connectomes, MD, FA, OD, ISOVF and ICVF were highly correlated with each other (*r* = 0.59 to 0.98). The hRSV based on SC had moderate correlations with other weighted hRSV (*r* = 0.23 to 0.35). Conversely, we saw that the hRSV has generally weak correlation with age, sex, WMV, GMV and atrophy, where the absolute values of correlations were mostly smaller than 0.1, see Figure 2b and 2c. We also see that the hRSV has weak-to-moderate correlation with all other graph measures (|*r*| : range = 0.07 to 0.33, mean = 0.23), see Figure 3.

### RSV, hRSV, assortativity, diffusion efficiency and global structural brain measures correlates significantly with *g*

We first examined the associations between *g* and RSV, hRSV, graph measures and global brain measures. The age, sex and MRI site corrected associations with graph measures are shown in Table 3.

**Table 3:** Associations between *g* and graph theoretical metrics based on six different structural network weights, RSV = Relative Strength Variability, 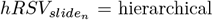 RSV with sliding window size *n*.

| Associations with $g$ | MD | FA | SC | OD | ISOVF | ICVF |
| --- | --- | --- | --- | --- | --- | --- |
| RSV | -0.0892 | **-0.0847** | 0.0122 | -0.0555 | -0.0525 | **-0.0770** |
| $hRSV_{slide_{21}}$ | -0.0678 | -0.0667 | -0.0467 | -0.0745 | -0.0562 | -0.0653 |
| $hRSV_{slide_4}$ | -0.0699 | -0.0717 | -0.0598 | **-0.0806** | **-0.0573** | -0.0697 |
| Mean Edge Weight | -0.0604 | 0.0433 | **0.1178** | -0.0029 | -0.0344 | 0.0396 |
| Clustering Coefficient | -0.0080 | -0.0025 | 0.0065 | -0.0061 | 0.0007 | -0.0043 |
| Assortativity | -0.0293 | -0.0231 | 0.0108 | 0.0209 | 0.0049 | -0.0050 |
| Strength Variance | -0.0042 | 0.0053 | 0.0247 | 0.0100 | 0.0020 | 0.0090 |
| Diffusion Efficiency | **0.0918** | 0.0781 | 0.0051 | 0.0678 | 0.0477 | 0.0740 |
| Routing Efficiency | 0.0096 | 0.0383 | 0.0179 | 0.0137 | 0.0008 | 0.0280 |

Higher RSV was significantly associated with lower *g* (*β*_*standardised*_ = −0.0892 to −0.0525, *p*_*FDR*_ = 8.46 × 10^*−*62^ to 4.1 × 10^*−*23^) for all the network weights except for SC (*β*_*standardised*_ = 0.0122, *p*_*FDR*_ = 0.0347), see Table 3 and Supplementary Materials B: Figure B.1. As for the hRSV, higher variability was significantly associated with lower *g* (*β*_*standardised*_ = −0.0806 to −0.0467, *p*_*FDR*_ = 3.54 × 10^*−*52^ to 1.20 × 10^*−*19^) for all the network weights, see Table 3 and Supplementary Materials B: Figure B.2. We saw that the variant with smaller window size (RSV_*slide*_4_) has minimally stronger associations with *g* than the one with larger window size (RSV_*slide*_21_). We also looked into the individual (and finer) sliding window measures calculated from low strength nodes to those calculated from high strength nodes, see Supplementary Materials B: Figure B.3. We see that ≥ 95% of the betas were negative, showing that the variability measure calculated across different ranges of node strengths were having robust associations with *g*, although the associations did not survive false discovery correction. On the other hand, we found that the associations between *g*, and the windowed variability measures computed from SC were rather inconsistent.

We also divided the nodes into four different tiers, echoing the work in binary hierarchical complexity (Blesa et al., 2021b; Smith et al., 2019). The associations with *g* were generally stronger for the windowed variability measures calculated from tier 1 and tier 3 nodes. For tier 1 nodes (i.e. nodes with top node strength), we also saw that the associations with *g* were more similar across different network weights when compared to nodes in other tiers, except for SC.

The diffusion efficiency had similar strength of associations to the RSV and hRSV with *g*, and was positively significantly associated with *g* (*β*_*standardised*_ = 0.0477 to 0.0918, *p*_*FDR*_ = 5.25 × 10^*−*65^ to 3.62 × 10^*−*19^) for all the network weights except for SC. Other graph measures had smaller effect sizes in terms of associations with *g* compared to the above two measures. The routing efficiency based on FA, SC, OD and ICVF were positively associated with *g* (*β*_*standardised*_ = 0.0137 to 0.0383, *p*_*FDR*_ = 6.95 × 10^*−*13^ to 0.0140). The assortativity scores based on MD, FA was negatively associated with *g* (*β*_*standardised*_ = −0.0293 to −0.0231, *p*_*FDR*_ = 9.06 × 10^*−*8^ to 1.57 × 10^*−*5^), while the one based on OD was positively associated with *g* (*β*_*standardised*_ = 0.0210, *p*_*FDR*_ = 2.30 × 10^*−*5^). The clustering coefficient scores were not significantly associated with *g* based on any of the network weights. Compared to the more conventional mean edge weight measure, we found that the RSV and hRSV has larger effect sizes in terms of associations with *g* for all the network weights (Mean Edge Weight: |*β*_*standardised*_| = 0.0029 to 0.0604, RSV, hRSV: |*β*_*standardised*_| = 0.0525 to 0.0892) except for SC, see Table 3 and Supplementary Materials B: Figure B.4 - B.8.

GMV, WMV, and brain atrophy had stronger associations with *g* than that of the RSV and hRSV (GMV: *β*_*standardised*_ = 0.2036, *p*_*FDR*_ = 5.57 × 10^*−*223^; WMV: *β*_*standardised*_ = 0.1568, *p*_*FDR*_ = 1.32 × 10^*−*133^; Brain atrophy: *β*_*standardised*_ = −0.1979, *p*_*FDR*_ = 1.91 × 10^*−*202^).

### Multivariate regression and incremental prediction of *g* by RSV, hRSV and other common graph measures

As above, we show correlations among graph measures (Tables 2a and 3), and then their bivariate associations with *g* (Table 3). We hypothesised that our novel measures would provide new information about cognitive differences beyond other network properties. We then formally quantified how far our novel measures explain unique variance in cognitive function beyond other established network measures. The results of the multivariable models, where mean edge weights, common graph theoretical metrics, and the RSV and hRSV were entered together to predict variance in *g*, are shown in Figure 4a. The clustering coefficient, node strength variance and routing efficiency are highly correlated with each other (*r* = 0.87 to 0.97, as stated in the above section), and, out of these three measures, routing efficiency had the most promising results in terms of associations with *g* (*β*_*standardised*_ = 0.0095 to 0.0383, *p*_*FDR*_ = 6.95 × 10^*−*13^ to 0.0915). Therefore, the clustering coefficient and node strength variance were not added to the multivariate models to reduce the severity of multicollinearity.

**Figure 4:**
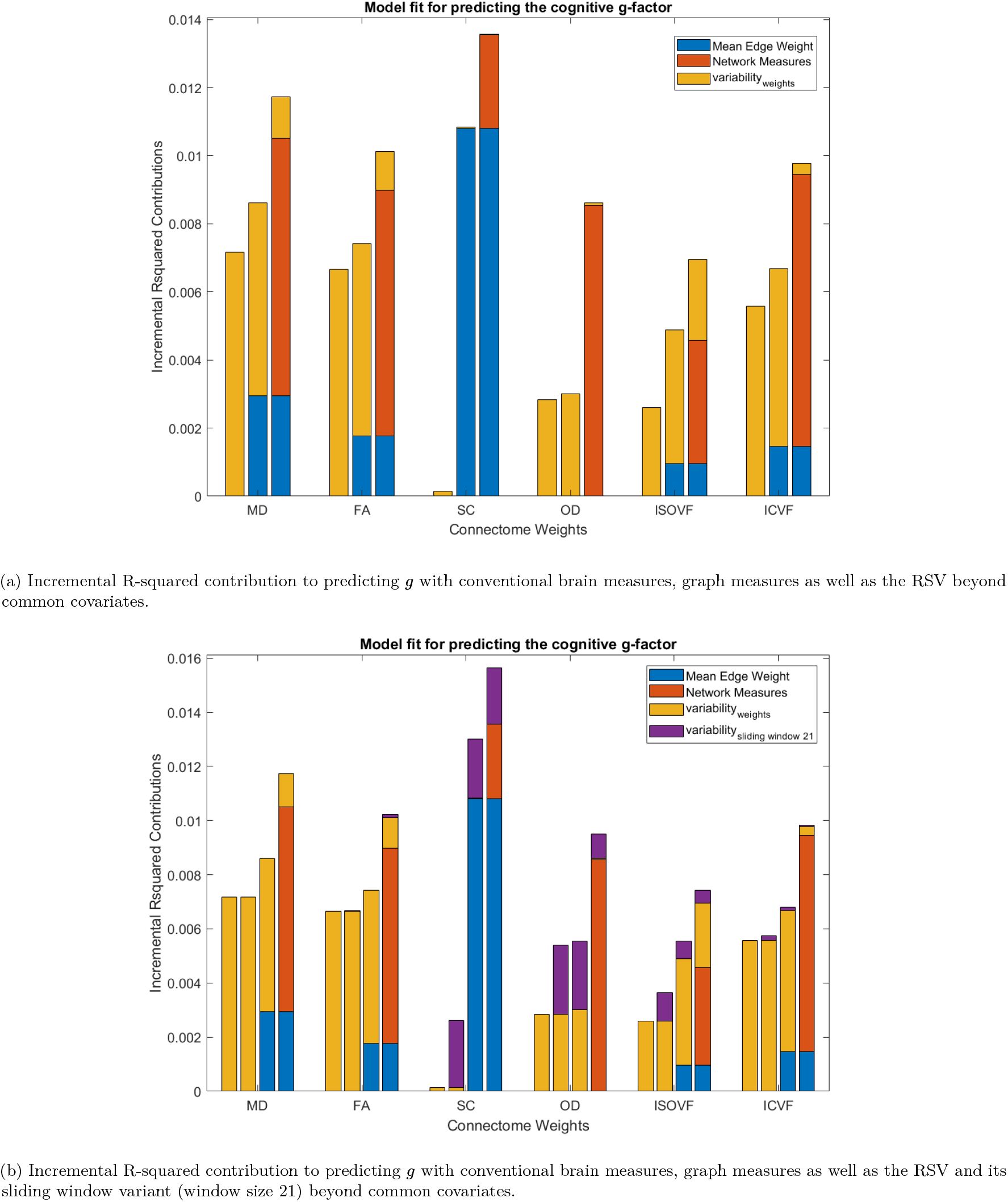
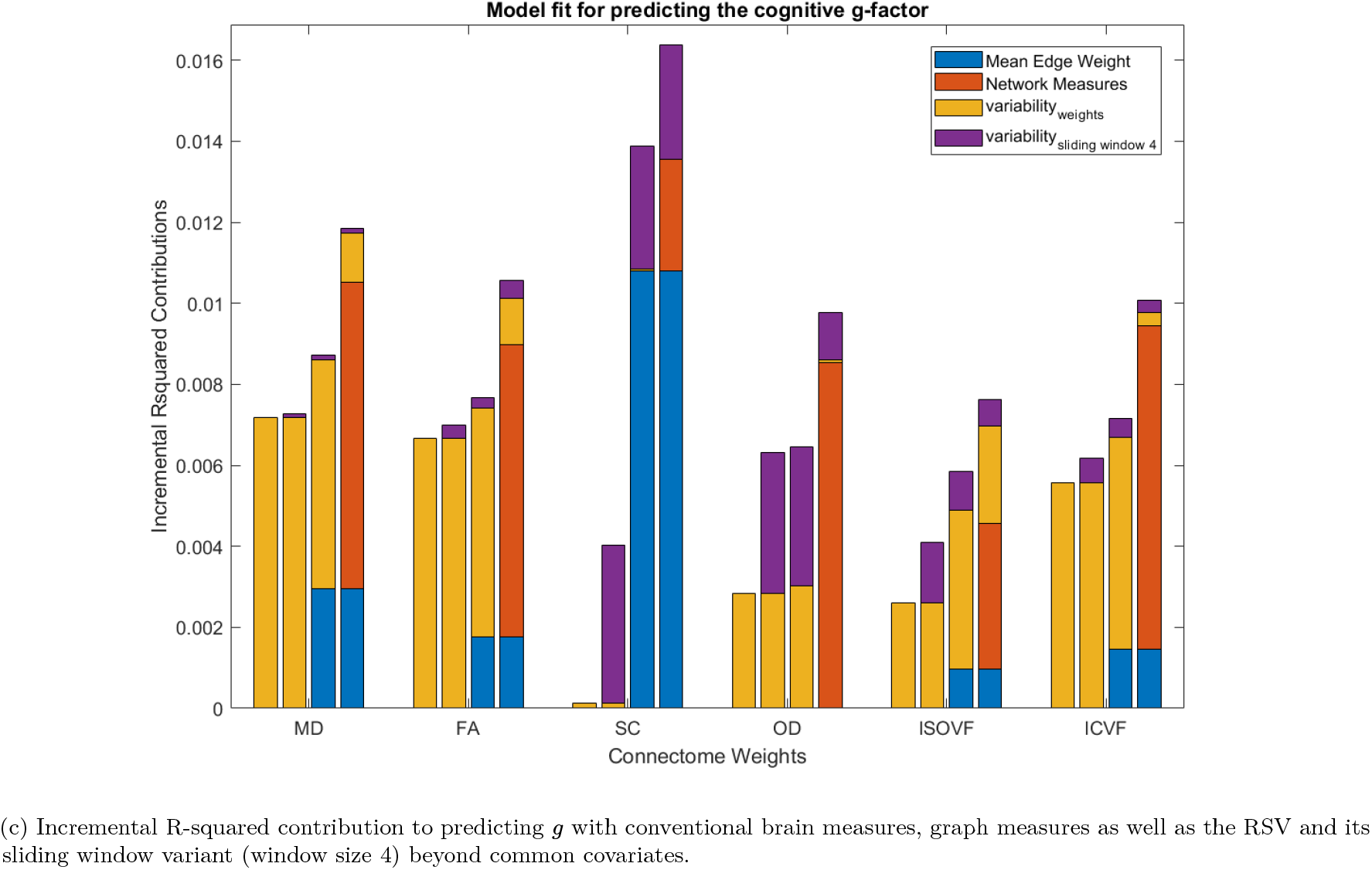
Incremental R-squared contribution to predicting *g* beyond common covariates. Note that the common covariates’ R-squared contribution to predicting *g* is 0.1448

Results showed that the RSV is incrementally significant for predicting *g* for all the network weights except for SC and OD (MD: *p*_*FDR*_ = 3.35 × 10^*−*6^, FA: *p*_*FDR*_ = 5.02 × 10^*−*6^, SC: *p*_*FDR*_ = 0.7055, OD: *p*_*FDR*_ = 0.2895, ISOVF: *p*_*FDR*_ = 5.36 × 10^*−*11^, ICVF: 0.0176), see Figure 4a. For SC, as we saw in the previous univariate analysis, the contribution was not significant and it was also not significant after adding other network measures. For OD, we found that the contribution by the RSV decreased to nearly zero after adding network measures to the multivariate regression. As for the sliding window variant, the incremental contribution of the 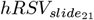 in predicting *g* was significant for SC, OD and ISOVF (MD: *p*_*FDR*_ = 0.5169, FA: *p*_*FDR*_ = 0.1793, SC: *p*_*FDR*_ = 1.07 × 10^*−*9^, OD: *p*_*FDR*_ = 1.03 × 10^*−*4^, ISOVF: *p*_*FDR*_ = 0.0049, ICVF: 0.3584), see Figure 4b. As for the 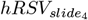 in predicting *g*, the incremental contribution was significant for all other network weights except for MD (MD: *p*_*FDR*_ = 0.1253, FA: *p*_*FDR*_ = 0.0044, SC: *p*_*FDR*_ = 5.90 × 10^*−*13^, OD: *p*_*FDR*_ = 5.77 × 10^*−*6^, ISOVF: *p*_*FDR*_ = 5.92 × 10^*−*4^, ICVF: 0.0190), see Figure 4c.

## Discussion

In the current study, we developed two novel measures to describe weighted brain structural connectivity: RSV, which is sensitive to network’s resilience to targeted attacks, and hRSV which we demonstrate as a weighted measure of statistical complexity of networks similar to the binary hierarchical complexity measure well-studied in structural connectomes (Blesa et al., 2021a; Smith et al., 2019). We computed the variability measures for brain structural connectomes derived from six different network weights, and assessed their relationship with age, sex, global brain measures as well as graph measures, in an exceptionally large sample of community-dwelling adults. We show in this large sample with wide age range that the routing efficiency, strength variance, assortativity and clustering coefficient decrease with advancing age while there is a subtle increase in RSV, and hRSV with advancing age. Individuals with lower RSV, hRSV and diffusion efficiency had subtly but significantly lower general cognitive function. Importantly, they demonstrated incremental significance beyond more conventional brain structural and other graph theory measures such as routing efficiency and clustering coefficient. The hRSV was also shown to have stronger association with *g* than the RSV for SC and OD. We also found that, whereas the mean weights in white matter that broadly represent directional coherence versus mean diffusivity (FA, ICVF vs MD and ISOVF respectively) are negatively correlated, the computations involved in RSV had a homogenising effect on the directions of associations with age, volumetric measures and *g* across weighting schemes.

RSV approximately measures the similarity between a node’s strength and strengths of its immediate neighbourhood, which is conceptually similar to measuring the local mixing patterns. A graph with higher RSV value partially indicates the existence of certain nodes with a majority of high-to-low strength connections, which may suggest more pronounced weak points in the network and therefore a higher vulnerability to targeted attack. Our results showed that the RSV and hRSV were significantly negatively associated with *g*, which lends some credence to such hypothesis, that brain network with higher vulnerability to targeted attack correlates with lower *g* in participant, mentioned earlier in the Introduction. For atlases with higher granularity, the higher-order standardised moments (e.g. skewness, kurtosis, etc.) may also be useful in describing the node relative strength distribution profile and further research will be needed to verify the clinical relevance of higher-order measures. Moreover, we also saw a weak but significant positive correlation between age and the RSV. Investigating the relationship between longitudinal change in the RSV and age-related cognitive decline will also be an interesting future research direction.

We showed that the hRSV is a weighted measure of statistical complexity which offers a distinct and complementary characterisation of relative strength connectivity patterns to RSV. Significantly, hRSV with a small window was comparatively weakly correlated with most demographic, structural and graph measures than RSV, yet was consistently as and often more strongly correlated with *g*. Interestingly, hRSV being negatively associated with *g* indicates that statistical complexity, particularly in high strength nodes, does not go hand-in-hand with general intelligence. This concurs with previous evidence which showed greater binarised connectome complexity in structural hubs in patients with lupus (Valdés Hernández et al., 2021), alongside a study showing that structural hubs had lower complexity, and so more regular connectivity patterns than random null models (Blesa et al., 2021b). The implication in this particular case then is that greater hRSV may be an indication of more random patterning, not necessarily more complex patterning.

When compared with other graph theory metrics, we found that the RSV and hRSV generally have stronger associations with *g* than the associations of other graph measures (i.e. clustering coefficients, strength variance, global routing efficiency and assortativity except for diffusion efficiency) with *g*. Previous studies have showed that measures of variance were more powerful in neuroimaging studies (Månsson et al., 2022), which is consistent with what we find in the current study. The global routing efficiency had comparatively small effect sizes with *g*. The global and local routing efficiencies were more commonly chosen as measures of brain network efficiencies in neuroimaging studies, possibly with the belief that signals were primarily propagated through shortest paths within the brain network. While we cannot comment on the truth of this for functional connectomes, this would categorically not be true for diffusion of water molecules forming the basis for structural connectomes. Using solely the efficiency defined by shortest paths did not take into account the possibility of signal propagation failure across longer-range brain connections. This may explain why the association between *g* and diffusion efficiency is stronger than that between *g* and routing efficiency. On the other hand, the diffusion efficiency was based on random walk on graphs and lacked the deterministic component. A brain network efficiency measure derived from biased random walks on graphs could be an interesting direction for future research.

Some limitations of the current study should be mentioned. UKB consists of healthier, wealthier, well-educated and older individuals (Fry et al., 2017), which may induce bias in the derivation of *g*. Network variables are known to be sensitive to the methods used for constructing structural connectomes (Qi et al., 2015), therefore results may differ with other processing setups. Moreover, we have chosen a particular atlas, the Desikan-Killany atlas, for constructing the brain network. Given that some graph metrics’ measurements are affected by atlas choice, the RSV could also be sensitive to parcellation schemes, and lead to different results. Further investigation will be needed to derive a harmonised RSV with improved robustness to choice of atlas.

## Conclusion

This study opens a new powerful avenue of exploration in weighted brain networks through the lens of relative node strength. We have used this framing to construct a measure of the degree of diversity in assortative mixing patterns in weighted networks, RSV, and the first statistical complexity measure for weighted networks. We demostrated that the RSV and hRSV have generally stronger associations with *g* than that of other graph theory metrics. Moreover, we also demonstrated that it contributes additional novel information to *g* predictions above other network measures, indicating cognitive decline is associated with more random hierarchical connectivity patterning in weighted structural connectomes.

## Acknowledgement and Funding Information

The research was conducted using the UK Biobank resource, with approved project number 10279. We are grateful to Prof Michelle Luciano for managing the team’s UK Biobank access. S. R. C. is supported by a Sir Henry Dale Fellowship, jointly funded by The Wellcome Trust and The Royal Society (221890/Z/20/Z), by the Milton Damerel Trust, and jointly by the UK BBSRC and ESRC (BB/W008793/1). Connectome generation, analyses, and H.W.Y., S. R. C., C. R. B., M. E. B., I. J. D., and E. M. T.-D. was supported by the US National Institutes of Health (National Institute on Aging; R01AG054628). S.R.C., I.J.D. and E.M.T.-D. are supported by the US National Institutes of Health (R01AG054628; U01AG083829).

## Notes

### Competing Interest Statement

The authors have declared no competing interest.

## References

Alfaro-Almagro, F., Jenkinson, M., Bangerter, N. K., Andersson, J. L., Griffanti, L., Douaud, G., Sotiropoulos, S. N., Jbabdi, S., Hernandez-Fernandez, M., Vallee, E., et al. (2018). Image processing and quality control for the first 10,000 brain imaging datasets from uk biobank. Neuroimage, 166:400–424.

Barbey, A. K., Colom, R., Solomon, J., Krueger, F., Forbes, C., and Grafman, J. (2012). An integrative architecture for general intelligence and executive function revealed by lesion mapping. Brain, 135(4):1154–1164.

Beaulieu, C. (2002). The basis of anisotropic water diffusion in the nervous system–a technical review. NMR in Biomedicine: An International Journal Devoted to the Development and Application of Magnetic Resonance In Vivo, 15(7–8):435–455.

Behrens, T. E., Berg, H. J., Jbabdi, S., Rushworth, M. F., and Woolrich, M. W. (2007). Probabilistic diffusion tractography with multiple fibre orientations: What can we gain? neuroimage, 34(1):144–155.

Behrens, T. E., Woolrich, M. W., Jenkinson, M., Johansen-Berg, H., Nunes, R. G., Clare, S., Matthews, P. M., Brady, J. M., and Smith, S. M. (2003). Characterization and propagation of uncertainty in diffusion-weighted mr imaging. Magnetic Resonance in Medicine: An Official Journal of the International Society for Magnetic Resonance in Medicine, 50(5):1077–1088.

Björck, Å. (1967). Solving linear least squares problems by gram-schmidt orthogonalization. BIT Numerical Mathematics, 7(1):1–21.

Blesa, M., Galdi, P., Cox, S., Sullivan, G., Stoye, D., Lamb, G., Quigley, A., Thrippleton, M., Escudero, J., Bastin, M., Smith, K., and Boardman, J. (2021a). Hierarchical complexity of the macro-scale neonatal brain. Cerebral Cortex, 31(4):2071–2084.

Blesa, M., Galdi, P., Cox, S. R., Sullivan, G., Stoye, D. Q., Lamb, G. J., Quigley, A. J., Thrippleton, M. J., Escudero, J., Bastin, M. E., et al. (2021b). Hierarchical complexity of the macro-scale neonatal brain. Cerebral Cortex, 31(4):2071–2084.

Buchanan, C. R. et al. (2020). The effect of network thresholding and weighting on structural brain networks in the UK Biobank. NeuroImage, page 116443.

Camilleri, J. A., Müller, V. I., Fox, P., Laird, A. R., Hoffstaedter, F., Kalenscher, T., and Eickhoff, S. B. (2018). Definition and characterization of an extended multiple-demand network. NeuroImage, 165:138–147.

Cox, S. R., Ritchie, S. J., Fawns-Ritchie, C., Tucker-Drob, E. M., and Deary, I. J. (2019). Structural brain imaging correlates of general intelligence in uk biobank. Intelligence, 76:101376.

Damoiseaux, J. S., Smith, S. M., Witter, M. P., Sanz-Arigita, E. J., Barkhof, F., Scheltens, P., Stam, C. J., Zarei, M., and Rombouts, S. A. (2009). White matter tract integrity in aging and alzheimer’s disease. Human brain mapping, 30(4):1051–1059.

Davenport, M. A. and Romberg, J. (2016). An overview of low-rank matrix recovery from incomplete observations. IEEE Journal of Selected Topics in Signal Processing, 10(4):608–622.

Deary, I. J., Corley, J., Gow, A. J., Harris, S. E., Houlihan, L. M., Marioni, R. E., Penke, L., Rafnsson, S. B., and Starr, J. M. (2009). Age-associated cognitive decline. British medical bulletin, 92(1):135–152.

Desikan, R. S. et al. (2006). An automated labeling system for subdividing the human cerebral cortex on MRI scans into gyral based regions of interest. Neuroimage, 31(3):968–980.

Eshkoor, S. A., Hamid, T. A., Mun, C. Y., and Ng, C. K. (2015). Mild cognitive impairment and its management in older people. Clinical interventions in aging, pages 687–693.

Fawns-Ritchie, C. and Deary, I. J. (2020). Reliability and validity of the UK Biobank cognitive tests. PloS one, 15(4):e0231627.

Foster, J. G., Foster, D. V., Grassberger, P., and Paczuski, M. (2010). Edge direction and the structure of networks. Proceedings of the National Academy of Sciences, 107(24):10815–10820.

Fry, A., Littlejohns, T. J., Sudlow, C., Doherty, N., Adamska, L., Sprosen, T., Collins, R., and Allen, N. E. (2017). Comparison of sociodemographic and health-related characteristics of uk biobank participants with those of the general population. American journal of epidemiology, 186(9):1026–1034.

Gläscher, J., Rudrauf, D., Colom, R., Paul, L. K., Tranel, D., Damasio, H., and Adolphs, R. (2010). Distributed neural system for general intelligence revealed by lesion mapping. Proceedings of the National Academy of Sciences, 107(10):4705–4709.

Goñi, J., Avena-Koenigsberger, A., Velez de Mendizabal, N., van den Heuvel, M. P., Betzel, R. F., and Sporns, O. (2013). Exploring the morphospace of communication efficiency in complex networks. PLoS One, 8(3):e58070.

Jones, D. K., Knösche, T. R., and Turner, R. (2013). White matter integrity, fiber count, and other fallacies: the do’s and don’ts of diffusion MRI. Neuroimage, 73:239–254.

Kennedy, K. M. and Raz, N. (2009). Aging white matter and cognition: differential effects of regional variations in diffusion properties on memory, executive functions, and speed. Neuropsychologia, 47(3):916–927.

Latora, V. and Marchiori, M. (2001). Efficient behavior of small-world networks. Physical review letters, 87(19):198701.

Li, X., Wang, Y., Wang, W., Huang, W., Chen, K., Xu, K., Zhang, J., Chen, Y., Li, H., Wei, D., et al. (2020). Age-related decline in the topological efficiency of the brain structural connectome and cognitive aging. Cerebral Cortex, 30(8):4651–4661.

Lyall, D. M., Cullen, B., Allerhand, M., Smith, D. J., Mackay, D., Evans, J., Anderson, J., Fawns-Ritchie, C., McIntosh, A. M., Deary, I. J., et al. (2016). Cognitive test scores in UK Biobank: data reduction in 480,416 participants and longitudinal stability in 20,346 participants. PloS one, 11(4):e0154222.

Madole, J. W., Buchanan, C. R., Rhemtulla, M., Ritchie, S. J., Bastin, M. E., Deary, I. J., Cox, S. R., and Tucker-Drob, E. M. (2023). Strong intercorrelations among global graph-theoretic indices of structural connectivity in the human brain. NeuroImage, 275:120160.

Månsson, K. N., Waschke, L., Manzouri, A., Furmark, T., Fischer, H., and Garrett, D. D. (2022). Moment-to-moment brain signal variability reliably predicts psychiatric treatment outcome. Biological Psychiatry, 91(7):658–666.

Mayo, C. D., Garcia-Barrera, M. A., Mazerolle, E. L., Ritchie, L. J., Fisk, J. D., Gawryluk, J. R., and Initiative, A. D. N. (2019). Relationship between dti metrics and cognitive function in alzheimer’s disease. Frontiers in aging neuroscience, 10:436.

Miller, K. L., Alfaro-Almagro, F., Bangerter, N. K., Thomas, D. L., Yacoub, E., Xu, J., Bartsch, A. J., Jbabdi, S., Sotiropoulos, S. N., Andersson, J. L., et al. (2016). Multimodal population brain imaging in the UK Biobank prospective epidemiological study. Nature neuroscience, 19(11):1523–1536.

Moodie, J. E., Harris, S. E., Harris, M. A., Buchanan, C. R., Davies, G., Taylor, A., Redmond, P., Liewald, D. C., Valdés Hernández, M. d. C., Shenkin, S., et al. (2024). General and specific patterns of cortical gene expression as spatial correlates of complex cognitive functioning. Human Brain Mapping, 45(4):e26641.

Newman, M. E. (2002). Assortative mixing in networks. Physical review letters, 89(20):208701.

Onnela, J.-P., Saramäki, J., Kertész, J., and Kaski, K. (2005). Intensity and coherence of motifs in weighted complex networks. Physical Review E, 71(6):065103.

Penke, L., Maniega, S. M., Murray, C., Gow, A. J., Hernández, M. C. V., Clayden, J. D., Starr, J. M., Wardlaw, J. M., Bastin, M. E., and Deary, I. J. (2010). A general factor of brain white matter integrity predicts information processing speed in healthy older people. Journal of Neuroscience, 30(22):7569–7574.

Qi, S., Meesters, S., Nicolay, K., ter Haar Romeny, B. M., and Ossenblok, P. (2015). The influence of construction methodology on structural brain network measures: A review. Journal of neuroscience methods, 253:170–182.

Smith, K. and Escudero, J. (2017). The complex hierarchical topology of eeg functional connectivity. Journal of Neuroscience Methods, 276:1–12.

Smith, K. M., Bastin, M. E., Cox, S. R., Valdés Hernández, M. C., Wiseman, S., Escudero, J., and Sudlow, C. (2019). Hierarchical complexity of the adult human structural connectome. Neuroimage, 191:205–215.

Smith, K. M. and Smith, J. P. (2024). Statistical complexity of heterogeneous geometric networks. PLOS Complex Systems, in press.

Smith, S. M., Jenkinson, M., Johansen-Berg, H., Rueckert, D., Nichols, T. E., Mackay, C. E., Watkins, K. E., Ciccarelli, O., Cader, M. Z., Matthews, P. M., et al. (2006). Tract-based spatial statistics: voxelwise analysis of multi-subject diffusion data. Neuroimage, 31(4):1487–1505.

Snijders, T. A. (1981). The degree variance: an index of graph heterogeneity. Social networks, 3(3):163–174.

Sporns, O., Tononi, G., and Kötter, R. (2005). The human connectome: a structural description of the human brain. PLoS computational biology, 1(4):e42.

Sudlow, C., Gallacher, J., Allen, N., Beral, V., Burton, P., Danesh, J., Downey, P., Elliott, P., Green, J., Landray, M., et al. (2015). Uk biobank: an open access resource for identifying the causes of a wide range of complex diseases of middle and old age. PLoS medicine, 12(3):e1001779.

Valdés Hernández, M. C., Smith, K. M., Bastin, M. E., Amft, E. N., Ralston, S. H., Wardlaw, J. M., and Wiseman, S. J. (2021). Brain network reorganisation and spatial lesion distribution in systemic lupus erythematosus. Lupus, 30(2):285–298.

Van Den Heuvel, M. P. and Sporns, O. (2011). Rich-club organization of the human connectome. Journal of Neuroscience, 31(44):15775–15786.

Wang, S.-P. and Pei, W.-J. (2008). First passage time of multiple brownian particles on networks with applications. Physica A: Statistical Mechanics and its Applications, 387(18):4699–4708.

Watts, D. J. and Strogatz, S. H. (1998). Collective dynamics of ‘small-world’networks. nature, 393(6684):440–442.

Yeung, H. W., Stolicyn, A., Buchanan, C. R., Tucker-Drob, E. M., Bastin, M. E., Luz, S., McIntosh, A. M., Whalley, H. C., Cox, S. R., and Smith, K. (2022). Predicting sex, age, general cognition and mental health with machine learning on brain structural connectomes. Human Brain Mapping.

Zhang, H., Schneider, T., Wheeler-Kingshott, C. A., and Alexander, D. C. (2012). NODDI: practical in vivo neurite orientation dispersion and density imaging of the human brain. Neuroimage, 61(4):1000–1016.

