## Supplementary Materials for "Relative strength variability measures for brain structural connectomes and their relationship with cognitive functioning"

March 7, 2025

#### A hRSV as a Weighted Statistical Complexity measure

A weighted measure of statistical complexity of networks should agree with the properties detailed for the binary case in (Smith and Smith, 2024). Namely, it should be 0 for regular graphs, tend to 0 for Erdös-Rényi graphs in the thermodynamic limit (as $n \to 0$) and be lower for more randomised models such as random geometric graphs and configuration models of networks. Here, we demonstrate that these properties are not all satisfied for RSV, but are all satisfied for hRSV with a $2\%n$ sliding window.

Firstly, regularity in a weighted network would entail the network had weights which were all equal. In this case, the relative strength of all nodes would be 1, for node $i$, the denominator being the mean of neighbourhood node strengths would be equal to the node strength of node $i$, so all relative node strengths would be equal and their variance would therefore be 0.

Now, we assess the behaviour of the metrics with respect to different weighted random models– Erdös-Rényi (ER) random graphs with weights $w_{ij} \sim U[0,1]$, random geometric graphs where weights are inverse distances $w_{ij} = exp(-d(i,j))$ for $d(i,j)$ the Euclidean distance between nodes $i$ and $j$, and random heterogeneous geometric graphs (Smith et al., 2019), where weights are $w_{ij} = exp(-d(i,j))(x_i + x_j)$ with $x_i \sim LN(0.5, 0.2)$ the parameter which controls degree heterogeneity in the network (here we use default parameters). Finally, we also study the configuration models of the heterogeneous geometric graphs. For this, the underlying binary networks are submitted to the standard configuration model procedure (links are disconnected leaving each node of degree $k_i$ with $k_i$ stubs which are then randomly paired across the network to create new links) and then the $k_i$ weights of node $i$ of degree $k_i$ are randomly assigned to the new randomised links of node $i$. The results for RSV and hRSV of these models with varying network densities ($d = 0.05, 0.1, \ldots, 0.95$), and network sizes ($n = 1000, 2000, \ldots, 10000$) are reported in Figure A.1.

We see that hRSV replicates the behaviour of the binary normalised hierarchical complexity measure (Smith and Smith, 2024), with hRSV of ER random graphs tending to 0 with increasing $n$ and heterogeneous geometric graphs having markedly larger values than their configuration models (random heterogeneous graphs) and random geometric graphs. While RSV also

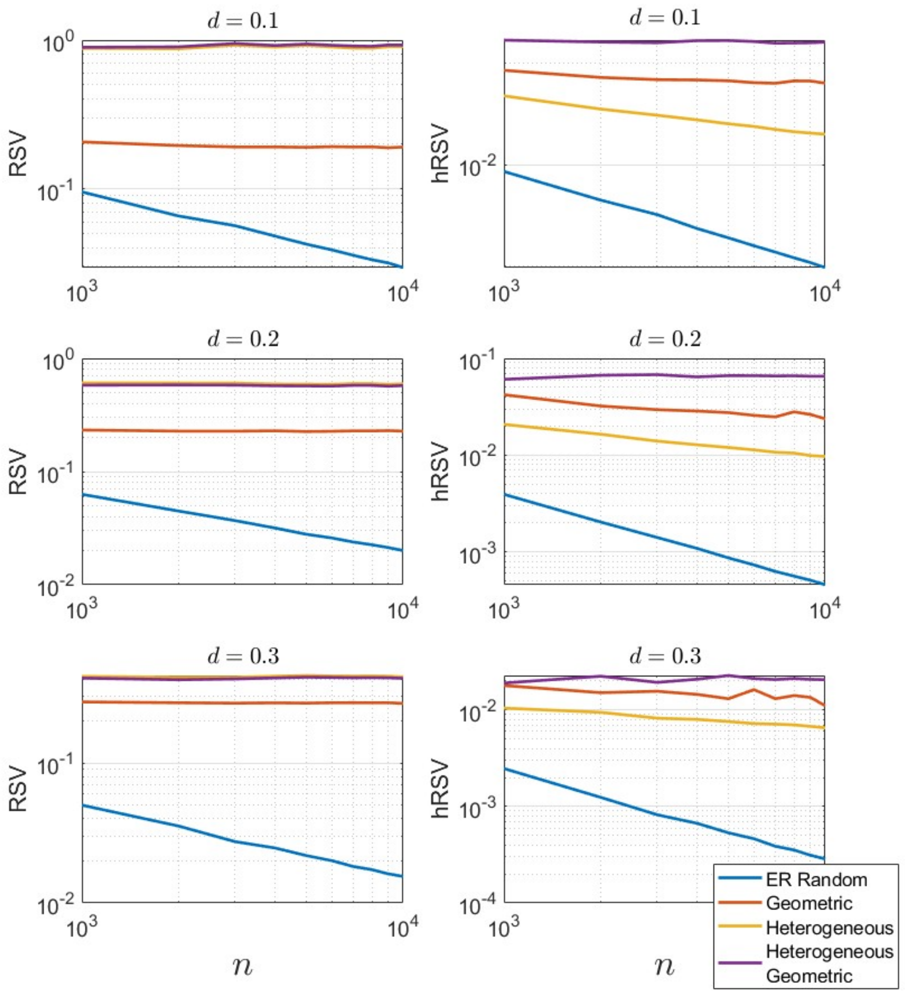


Figure A.1: The behaviour of Relative Strength Variability (RSV) and hierarchical RSV (hRSV) with respect to random weighted network models with varying $n$ (on the $x$-axis) and $d$ (as indicated in plot titles). Models shown are Erdös-Rényi random graphs (ER random), random geometric graphs (Geometric), random heterogeneous graphs (heterogeneous) and random heterogeneous geometric graphs (heterogeneous geometric), as in the legend.

shows tending to 0 for ER random graphs, it is clear that it does not distinguish well between heterogeneous geometric graphs and their configuration models. Taken together, we can propose hRSV as a measure of statistical complexity for weighted networks, but not RSV.

Notably, both RSV and hRSV appear to be well normalised with respect to network size for heterogeneous geometric graphs. This is important because these graphs are good models of real-world networks which indicates the measures can be used straightforwardly to compare real-world networks of different sizes.

The behaviour of the metrics with respect to network degree heterogeneity was then also explored. Here, the paremeter $\sigma$ in the heterogeneous geometric graphs is varied. Values are computed 100 times for each $\sigma = 0.05, 0.1, \ldots, 1$, density $d = 0.3$ (matching the structural connectomes), and sliding windows $w = 5, 10, 25, 50, 125, 250, 500$. Results are shown in Figure A.1 for $n = 1000$ and $n = 85$. Here, we vary the window size for hRSV from $n/2$ to $\sim 2\%n$, with all choices shown in the legends.

Again, for small $w$ we replicate the behaviour seen for normalised hierarchical complexity in (Smith and Smith, 2024) with the highest values achieved for graphs with $\sigma \in [0.2, 0.4]$. Interestingly, for RSV the values continue to increase with increasing $\sigma$, while for intermediate $w$ the prevailing behaviour is a levelling off with large $\sigma$. This motivates the selection of

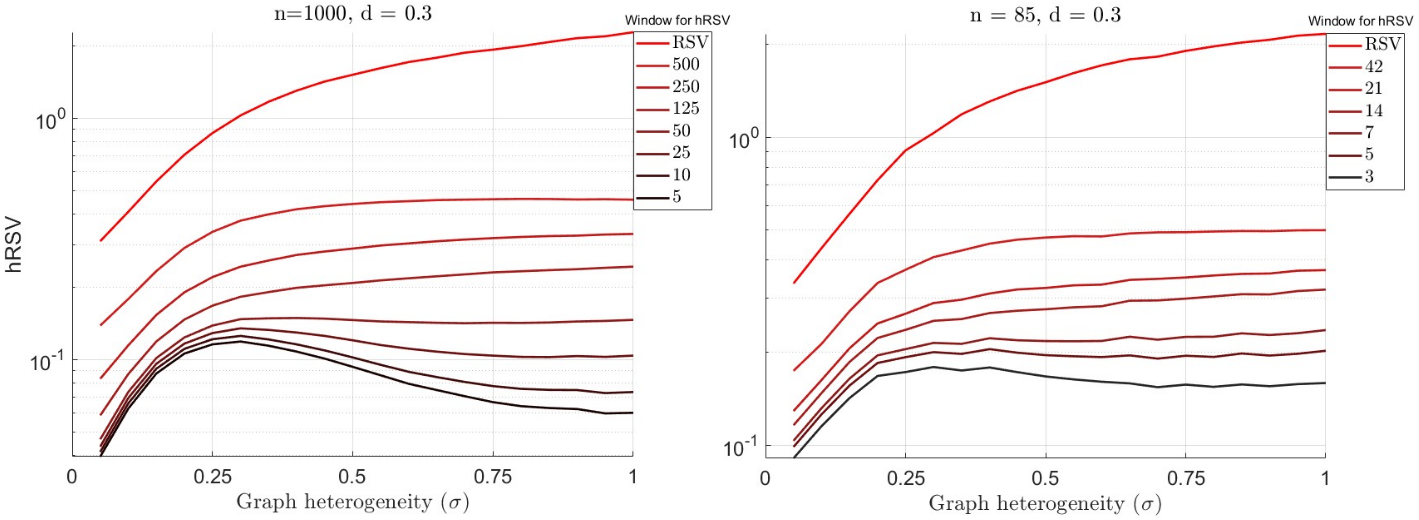


Figure A.2: The behaviour of hierarchical Relative Strength Variability (hRSV) measures with respect to geometric network degree heterogeneity.

a suitably small $w$ for the real world applications described in the results, to best distinguish the hRSV measure from the standard RSV.

#### A.1 Figures showing correlations among the graph measures for 6 different network weights

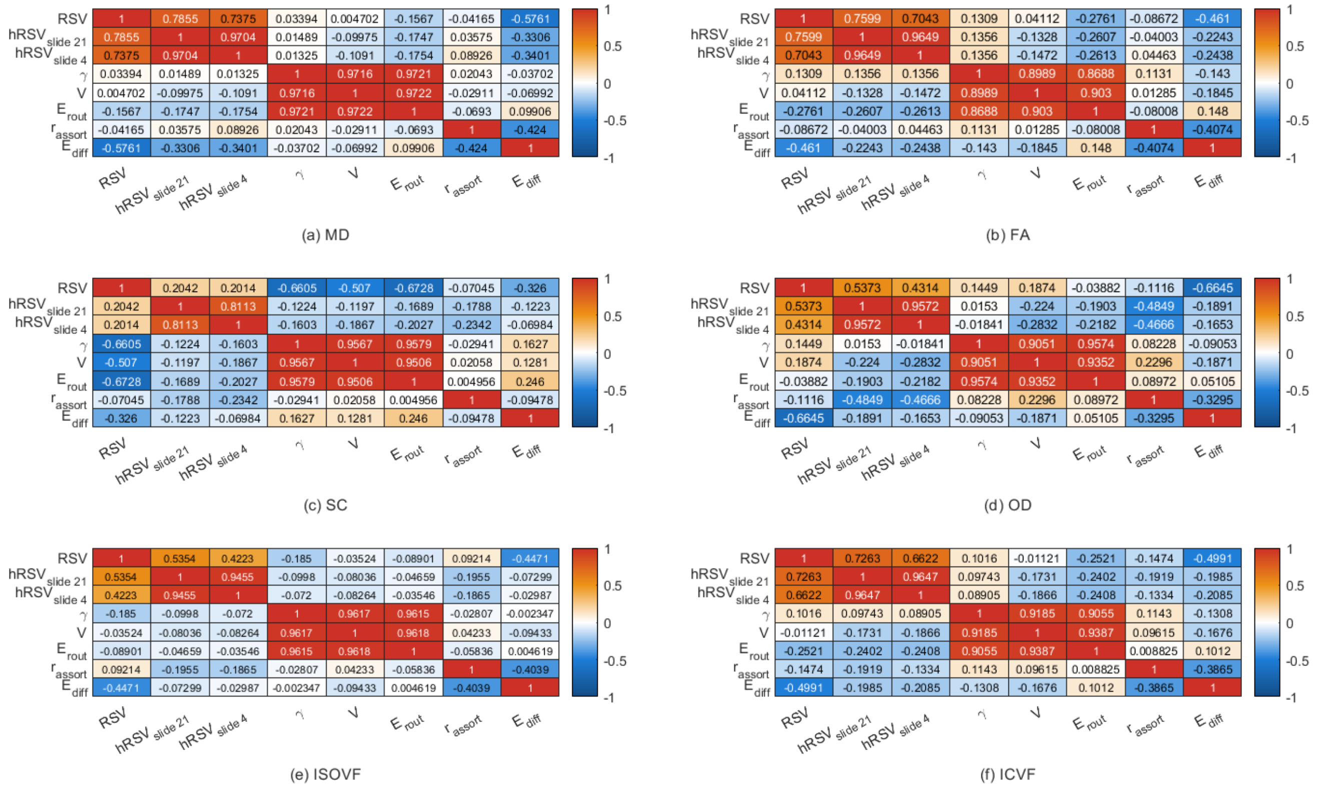


Figure A.3: Heatmaps showing correlation among the five chosen graph metrics based on different connectome weights. RSV = Relative Strength Variability, hRSV$_{slide_n}$ = Relative Strength Variability sliding window variant where $n$ is the window size, $\gamma$ = Clustering Coefficient, $r_{assort}$ = assortativity, $V$ = node Strength Variance, $E_{diff}$ = Global Diffusion Efficiency, $E_{rout}$ = Global Routing Efficiency

### B Figures showing age- and sex-corrected associations between $g$-factor and graph measures

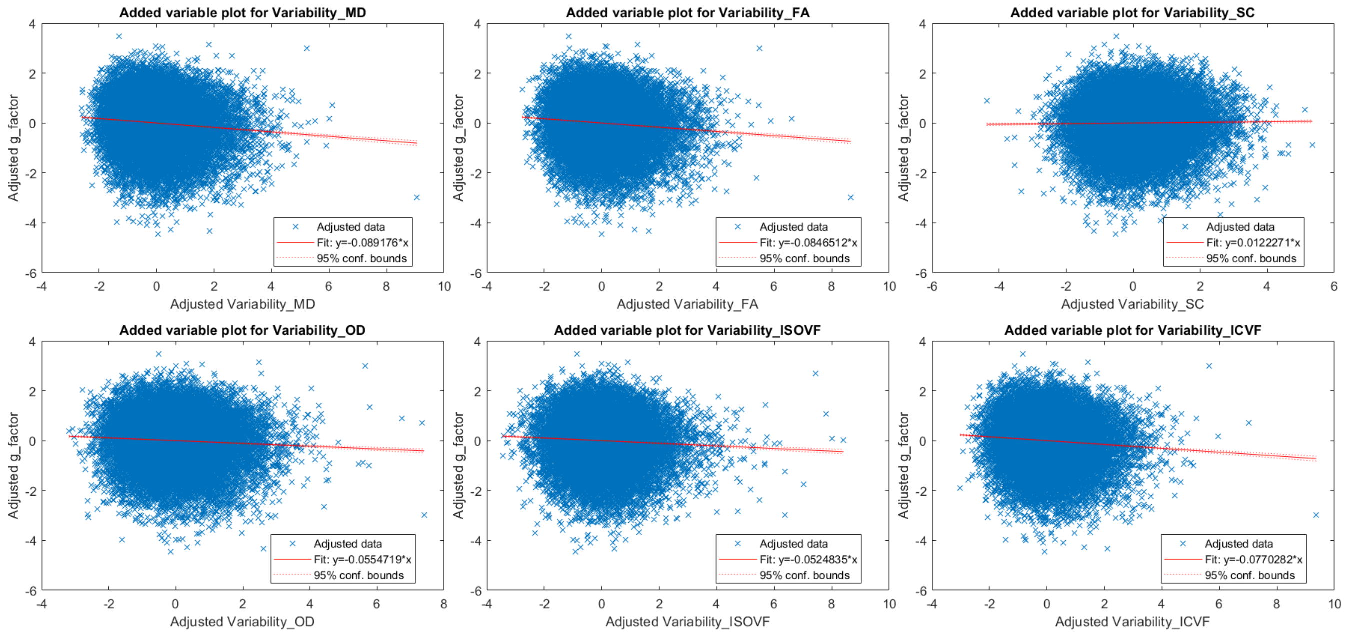


Figure B.1: Scatter plots showing the associations between the variability measure and the $g$-factor, adjusted for common covariates, for each of the six network weights.

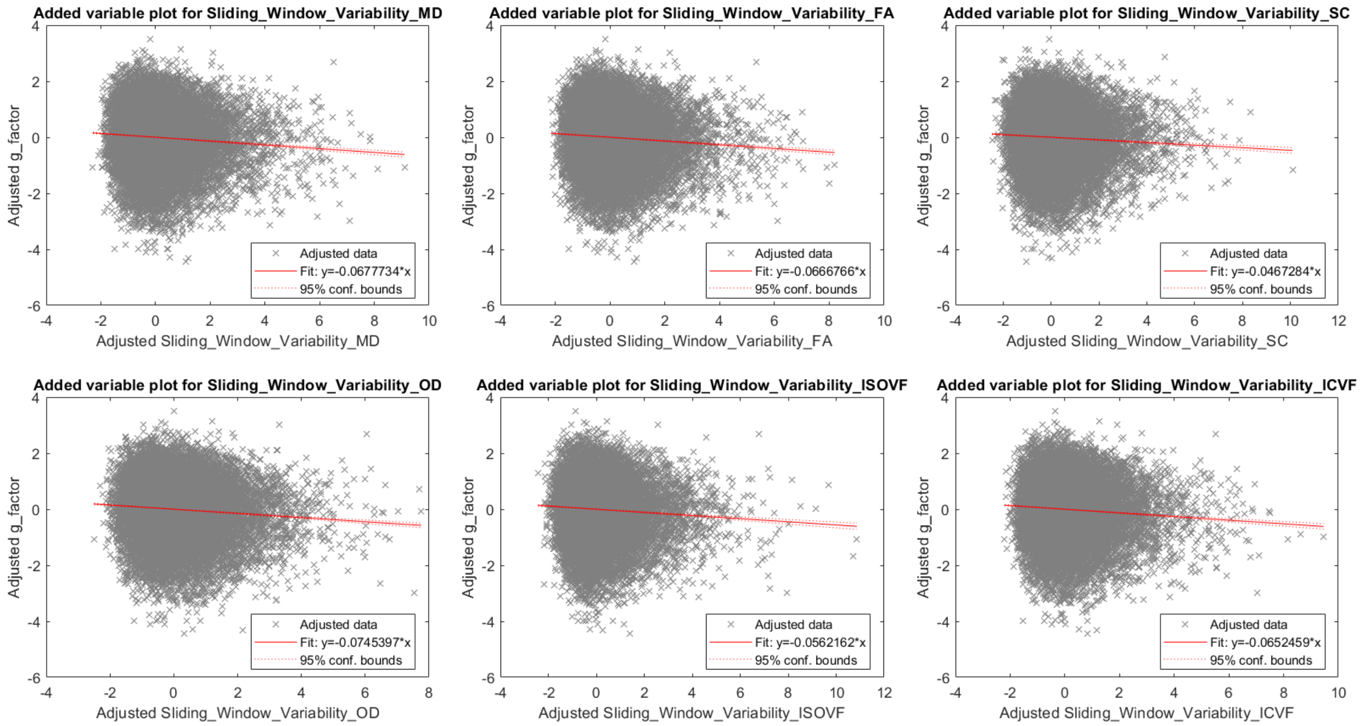


Figure B.2: Scatter plots showing the associations between the fine sliding window variability measures and the $g$-factor, adjusted for common covariates, for each of the six network weights.

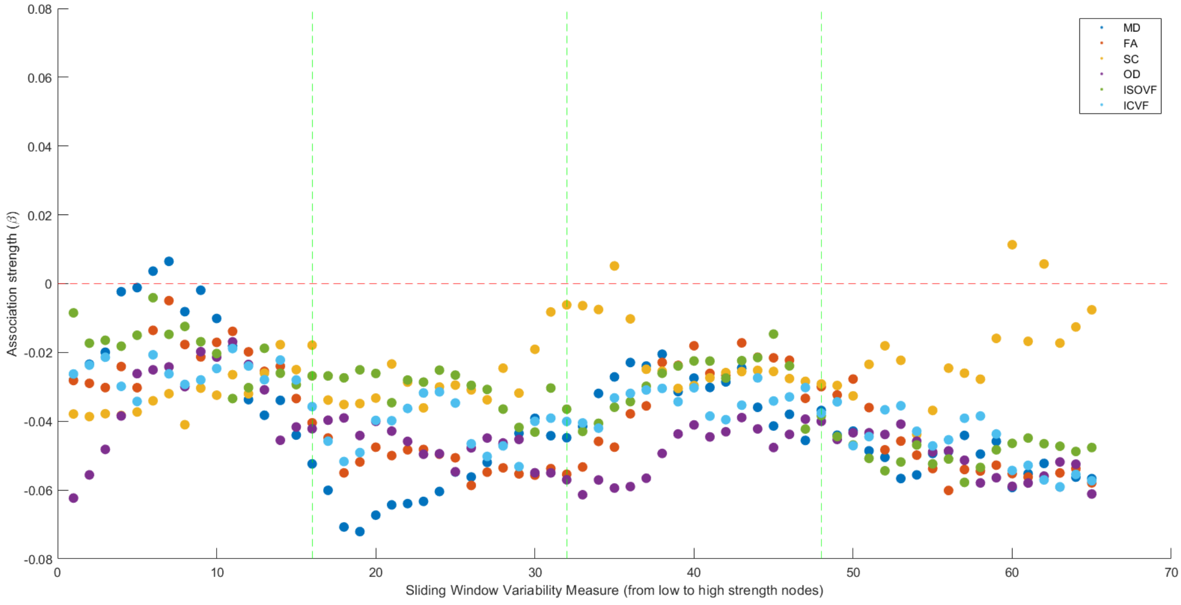


Figure B.3: Scatter plots showing the associations between the fine sliding window variability measures and the *g*-factor, adjusted for common covariates, for each of the six network weights.

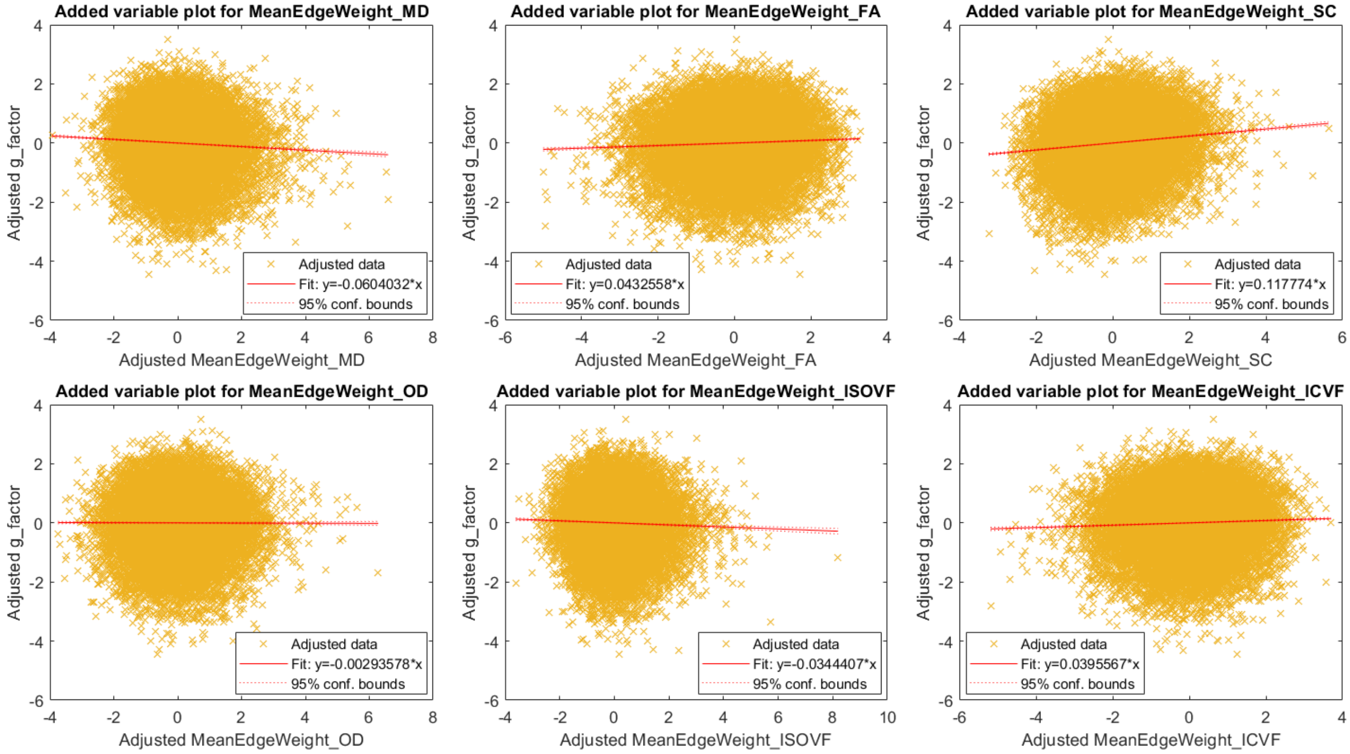


Figure B.4: Scatter plots showing the associations between mean edge weight and the *g*-factor, adjusted for common covariates, for each of the six network weights.

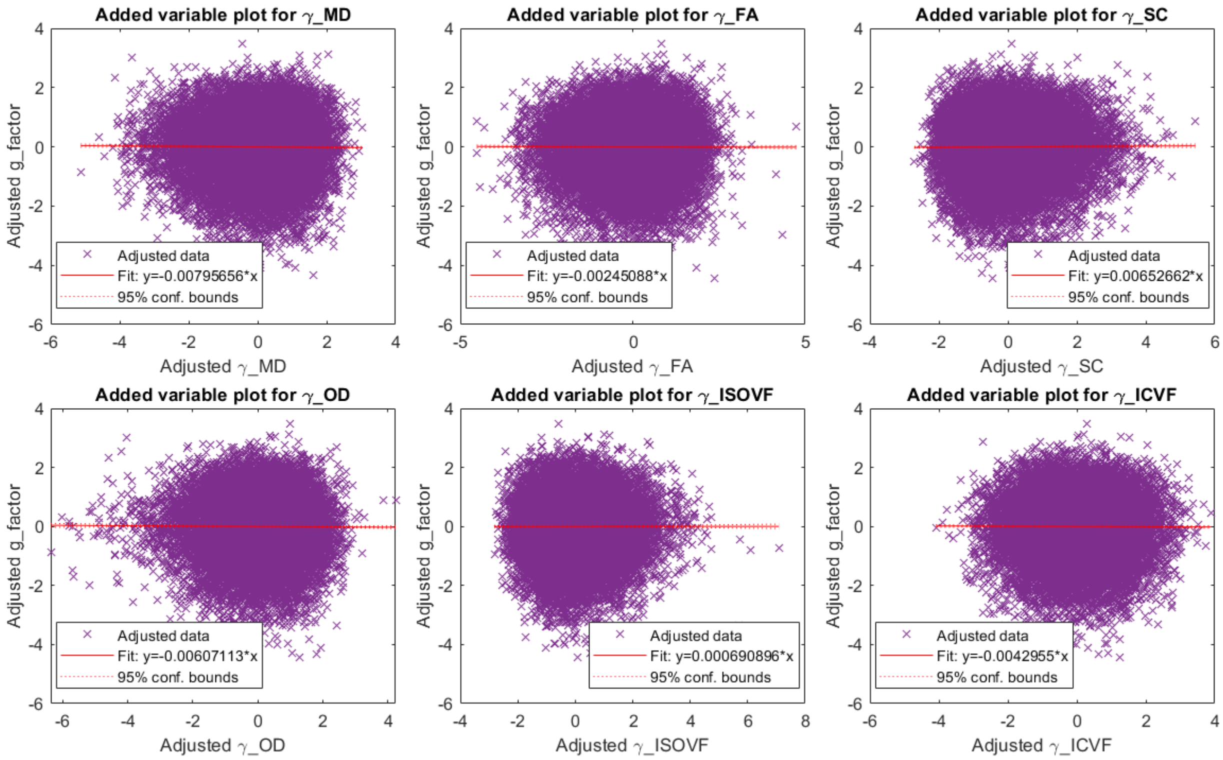


Figure B.5: Scatter plots showing the associations between clustering coefficients and the $g$-factor, adjusted for common covariates, for each of the six network weights.

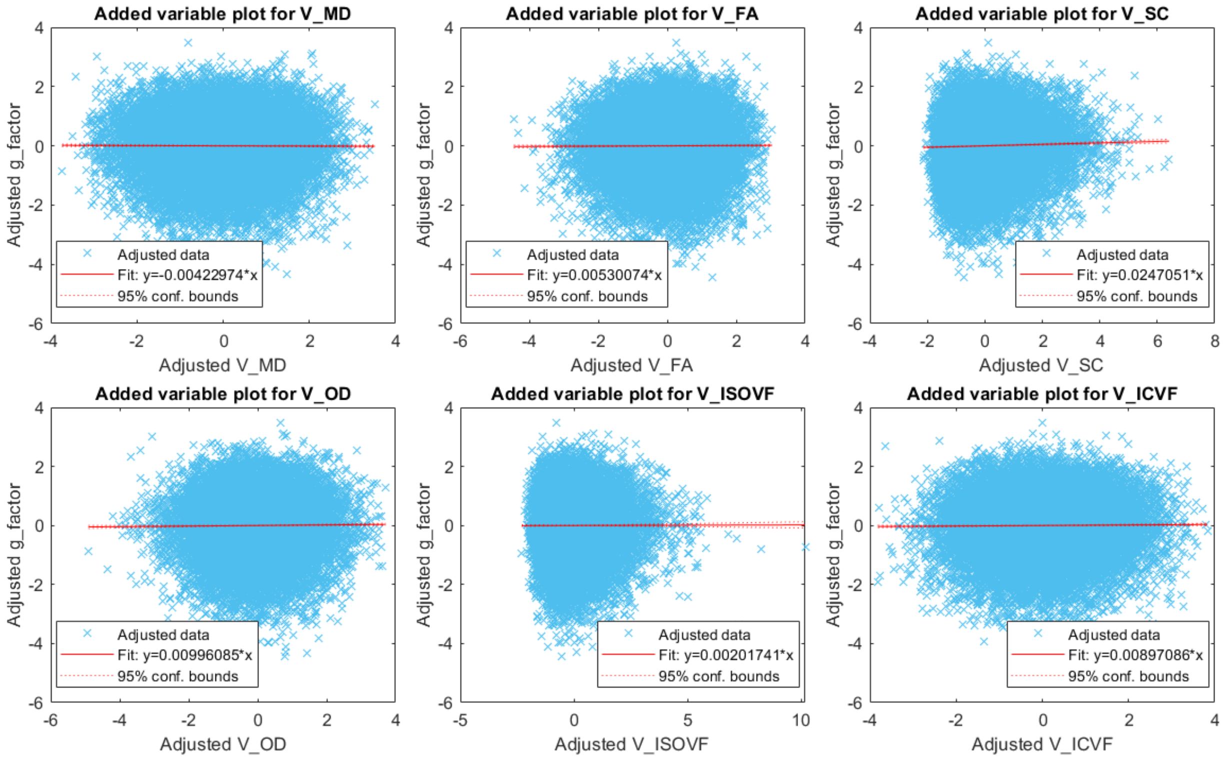


Figure B.6: Scatter plots showing the associations between strength variance and the $g$-factor, adjusted for common covariates, for each of the six network weights.

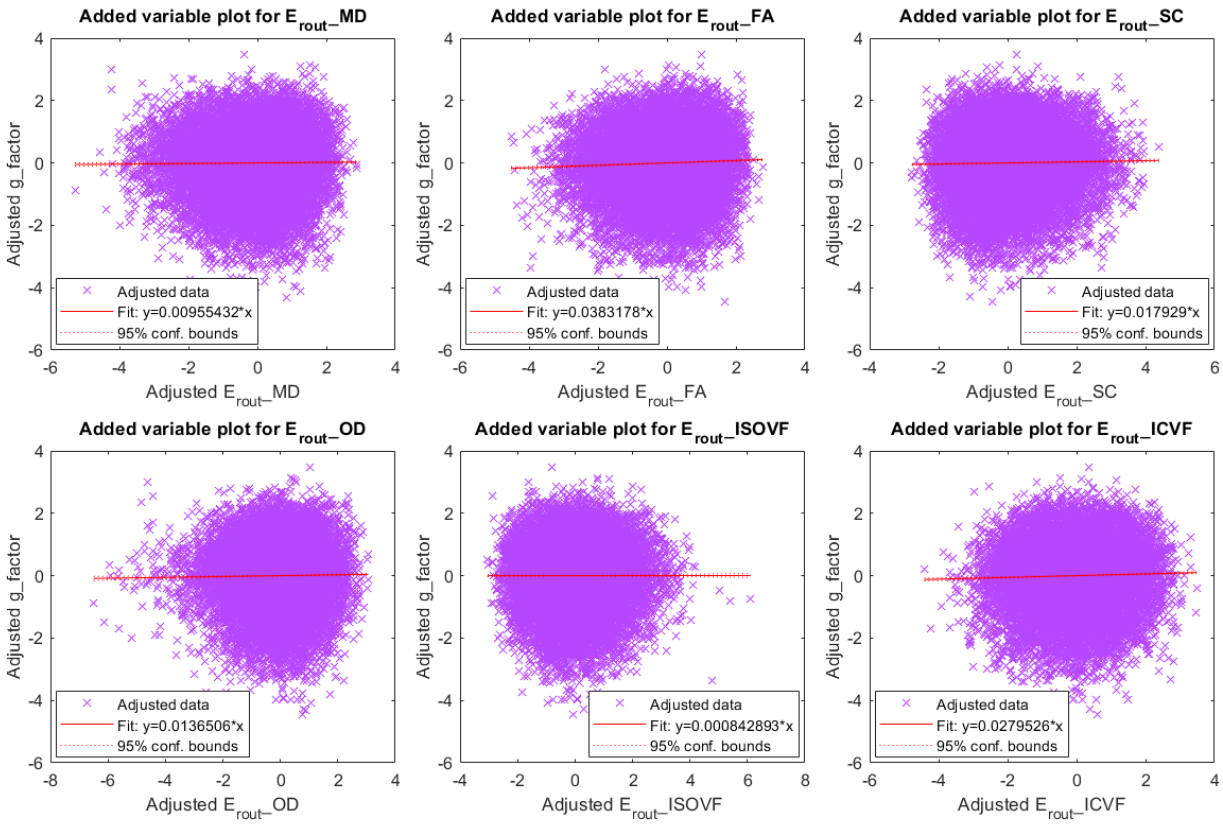


Figure B.7: Scatter plots showing the associations between routing efficiency and the $g$-factor, adjusted for common covariates, for each of the six network weights.

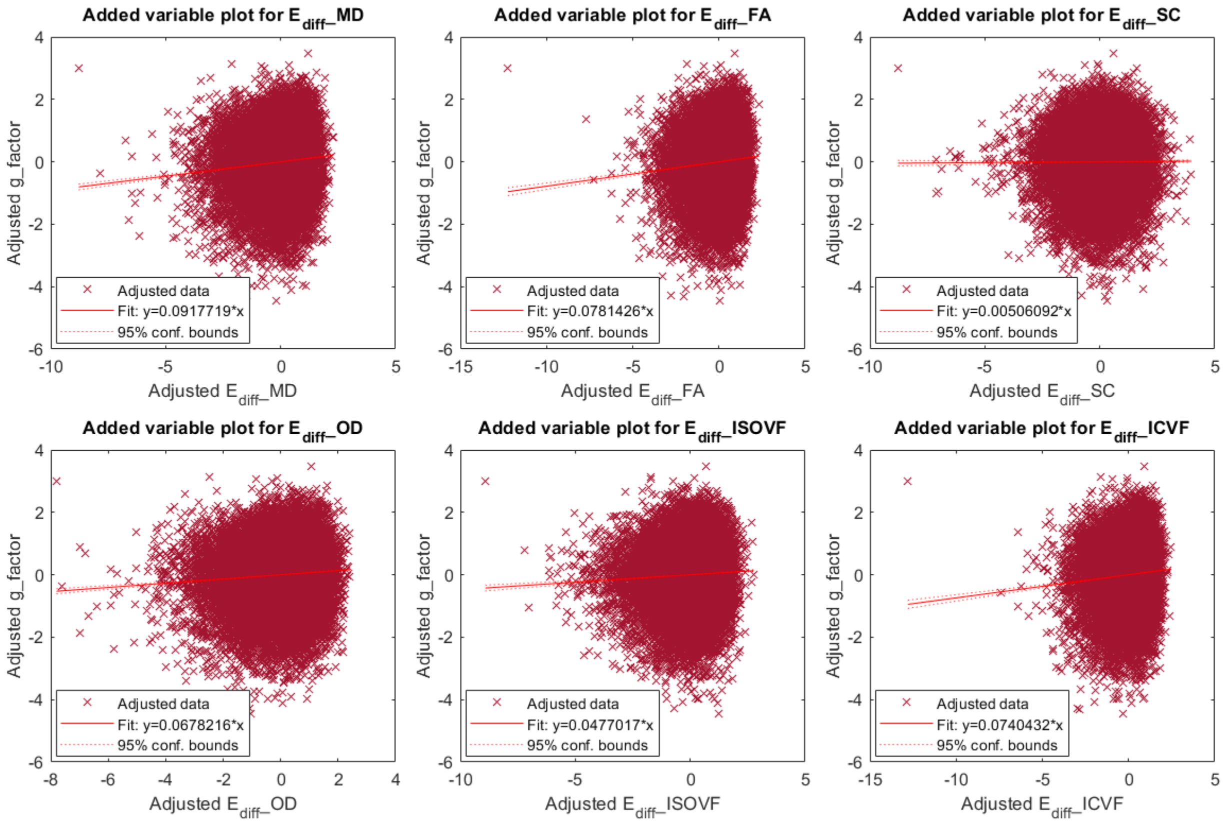


Figure B.8: Scatter plots showing the associations between the diffusion efficiency and the $g$-factor, adjusted for common covariates, for each of the six network weights.

### References

Smith, K. M., Bastin, M. E., Cox, S. R., Valdés Hernández, M. C., Wiseman, S., Escudero, J., and Sudlow, C. (2019). Hierarchical complexity of the adult human structural connectome. *Neuroimage*, 191:205–215.

Smith, K. M. and Smith, J. P. (2024). Statistical complexity of heterogeneous geometric networks. *PLOS Complex Systems*, in press.
